# Drinking water treatment drastically modifies multi-kingdom water communities, but clinically relevant opportunistic bacterial pathogens persist

**DOI:** 10.64898/2026.09.16.751965

**Authors:** Pierre Foucault, Ana Elena Pérez-Cobas, Madalina Ababii, Jesus Marin-Miret, Sebastien Wurtzer, Laurent Moulin, Carmen Buchrieser, Laura Gomez-Valero

## Abstract

Controlling pathogenic microorganisms during water treatment processes is a major public health concern. However, the impact of different water sources and treatment processes on the drinking water microbiome and opportunistic bacterial pathogens remains poorly understood. Here, we report the analyses of two water systems: a river and groundwater source, and their corresponding treated waters, sampled bi-weekly for 6 months. Microbial communities were analysed through a metabarcoding and dPCR workflow coupling universal prokaryotic and eukaryotic primers with genus-targeting primers for *Legionella* and *Pseudomonas* to achieve a deep taxonomic resolution. We show that microbial communities in groundwater were more diverse than those in river water and that water treatment affected α-diversity. Water treatment had a stronger influence on community composition than water system or seasonal variation. Despite lower bacterial loads, the relative abundance of *Legionella*, *Pseudomonas*, and non-tuberculous *Mycobacteria* (NTM) remained stable or even increased after water treatment, suggesting that conventional treatment processes promote enrichment of specific taxa. Excitingly, the genus-targeting analyses revealed a remarkable and previously uncharacterized diversity of *Legionella* species. *L. pneumophila*, pathogenic *Pseudomonas* clades and *P. aeruginosa* were detected in all environments, emphasizing their ability to adapt to diverse ecological conditions. Analysis of the eukaryotic fraction emphasized the persistence and emergence of eukaryotic hosts associated with these waterborne opportunistic pathogens. Our findings offer important insights into how water source characteristics and treatment processes influence the composition of the drinking water microbiome, while the high taxonomic resolution achieved for waterborne opportunistic bacterial pathogens underscores their ecological adaptability.

## Introduction

Water treatment has been a major advancement in public health, providing access to safe drinking water and drastically reducing diseases related to waterborne pathogens such as diarrhoea, cholera, dysentery and typhoid fever. The efficiency of this water treatment has been traditionally assessed by measuring faecal indicator bacteria. Although these bacteria are generally not harmful themselves, they serve as indicators of a potential faecal contamination^1^. However, these controls do not detect opportunistic waterborne pathogens. Indeed, while water treatment has been successful in removing faecal pathogens, it has shifted the waterborne diseases from gastrointestinal illnesses (such as cholera and typhoid fever) to primarily respiratory and cutaneous diseases caused by microorganisms that can persist despite treatment^2^. These organisms are generally known as opportunistic premise plumbing pathogens^3^ or opportunistic waterborne pathogens and they are transmitted primarily through inhalation of aerosols or direct contact with water, rather than by ingestion. In this group, most waterborne diseases and related deaths can be attributed to three bacterial genera: *Legionella*, *Pseudomonas*, and non-tuberculous *Mycobacteria* (NTM)^4^. These pathogens share the ability to form biofilms and interact with protists such as amoebae and ciliates, allowing them to survive harsh conditions, such as the presence of chlorine. These pathogens are considered “opportunistic” because they primarily cause disease in individuals with underlying risk factors, such as advanced age, cancer, or immunodeficiency, a growing proportion of the population^5^.

Recent scientific evidence also highlights the link between climate change, manifested by rising water temperatures and increased precipitation, and the growing environmental abundance of waterborne pathogens such as *Legionella*. Indeed, infections caused by *Legionella*, *Pseudomonas*, and non-tuberculous *Mycobacteria* (NTM) are steadily increasing worldwide^6–9^. This trend also extends to amoebae, which serve as hosts for these pathogens, as evidenced by reports linking environmental parameters, susceptible to alteration by climatic change, to amoebal abundance^10^. Thus, waterborne opportunistic pathogens represent an emerging infectious disease threat that warrants urgent attention, as standard coliform monitoring and even mandatory Heterotrophic Plate Count tests to detect re-growth are merely indirect indicators that lack species specificity^11^.

Although the implementation of advanced metagenomic surveys to study drinking water over the past years has provided a more comprehensive view of the drinking water microbiome, most studies analysing drinking water treatment plants do not include the water sources supplying the plant^12^. Furthermore, only few studies analyse both, the prokaryotic and eukaryotic fraction of the microbiome^13,14^, even though eukaryotic hosts are essential for the survival of several bacterial pathogens. Finally, the identification of bacteria rarely reaches sufficient taxonomic resolution to identify organisms beyond the genus level^7^, which limits the ability to accurately identify pathogenic species. In our study, we sought to provide a more comprehensive characterization of the water microbiome and improve our understanding of strategies to prevent and control bacterial infections caused by waterborne opportunistic pathogens. Thus, we analysed the prokaryotic and eukaryotic communities from two different water sources within the Paris drinking water network, a river and a mix of connected underground sources. These two different sources and the treated water leaving their respective treatment plants were sampled biweekly from February to August 2023 and the prokaryotic and eukaryotic fractions of the water microbiome were analysed. These were combined with an in-depth analysis of selected bacterial waterborne pathogens to characterize their frequency, abundance, and diversity at high taxonomic resolution. We propose that different raw water sources harbour unique microbial communities, and that after treatment the resulting communities are shaped by both the type of water source and the specific treatment processes applied. We further reveal that the community composition of both systems converges after treatment, particularly with respect to disinfection-resistant microorganisms, due to their shared capacity to survive treatment.

## Materials and methods

### Water sample collection and processing

Samples were collected bi-weekly (February to August 2023) at the entry (raw water) and exit (treated water) of two water treatment plants (Joinville and Haÿ-les-Roses) alimenting Paris’ drinking water. Joinville is fed from the Marne River (river water, RW), and releases treated river water (TRW). Haÿ-les-Roses is alimented from various underground sources transported through the Vanne aqueduct (groundwater, GW) and releases treated groundwater (TGW). For detailed treatment processes see supplementary materials. Water temperature, total organic carbon (COT), nitrates (NO ions), nitrites (NO_2_ ions), orthophosphates (PO ³ ions), pH, free chlorine (free Cl ions), iron (Fe^2+^ ions) and water conductivity were measured (**Table S1**). 1L water bottles (with 20 mg of NaS_2_O_3_ to neutralize residual chlorine in TW samples) were stored at 4 °C and transported to the laboratory within 12 hours. 1L of RW and 3L of TW were filtered through a sterile polycarbonate membrane filter (0.4 μm, Millipore®) in a filtration manifold, with a filtration blank included.

### Nucleic acid extraction

A combined chemical (800 µL of Trizol) and mechanical (Fisherbrand™ Bead Mill 24 homogenizer, 10 minutes, 3.1 m/s) lysis approach was performed at room temperature. The tubes were centrifuged, the supernatant was collected and nucleic acid purification was carried out using a QIAsymphony automated extractor (QIAGEN) with the DSP Virus/Pathogen Kit (#937055), following the manufacturer’s instructions. The extracted nucleic acids were eluted in a final volume of 50 µL of elution buffer and stored at 4 °C.

### Absolute quantification

Digital PCR (dPCR) was performed using the QIAcuity instrument (QIAGEN) with the QIAcuity PCR Probe Master Mix Kit (#250098) on the Nanoplate 26k 24-well plate (#250102) following the manufacturer’s instructions. Primers and probes are listed in **Table S2**. The cycling protocol and data processing is detailed in supplementary materials.

### Amplicon PCR, Illumina MiSeq library preparation and sequencing

Universal rRNA encoding gene primer sets were used for prokaryotes (515Fb/806Rb)^15,16^ and eukaryotes (1427F/1616R)^17^ and genus-targeting primer sets were used for *Legionella* (Lgsp17F/28R)^18^ and *Pseudomonas* (Pse464F/665R)^19^. The cycling protocol is detailed in supplementary materials. Libraries were prepared following the manufacturer’s instructions using the Nextera XT DNA Library Preparation Kit (Illumina, #FC-131-102) and send to sequencing on the iGenSeq platform (Illumina MiSeq, 2×250 bp; Paris, France). Amplicon sequence pre-processing is detailed in supplementary materials.

### Curated taxonomic annotation of *Legionella* and *Pseudomonas* ASVs

Curated in-house strain-level databases were created for Legionellaceae and Pseudomonaceae.16S rRNA sequences were downloaded from four existing curated databases: LPSN^20^, NCBI Bacterial 16S Ribosomal RNA RefSeq Targeted Loci Project, MIMt 16S-M2c-24-10^21^, and GTDB-r226^22^. Representative sequences of outgroup taxa from GTDB-r226^22^ were added. Strain names were manually checked and corrected for consistency across databases. Sequences were trimmed, quality-filtered and strain-level dereplicated, yielding 1235 Legionellaceae and 734 Pseudomonaceae strain-level sequences. The pipeline is detailed in the in supplementary materials. ASV_L_ and ASV_P_ sequences were blasted against their respective database (blastn^23^ v2.15.0, evalue <= 1e-10). Multiple annotations and a consensus were reported in case of equal best BIT scores (**Table S7, S8, S9**). Based on intra- and inter-species pairwise comparisons (supplementary materials), ASVs_L_ were classified according to sequence identity of =>99% as “very closely related”, of 97-99% as “closely related” or “distantly related” (<97%) to a known Legionellaceae species (**Fig. S1**). ASVs_P_ were classified as “very closely related” (=>99%), “closely related” (95-99%) or “distantly related” (<95%) to a known Pseudomonaceae species (**Fig. S2**).

### Phylogenetic analysis

The most abundant ASVs_L_ and ASVs_P_ were defined as the 10 ASVs with the highest average relative abundance by site. Their sequences, together with the species-level dereplicated sequences (or clade representative species for Pseudomonaceae^24,25^; **Table S10**) from their respective databases, were aligned using SINA^26^ (v1.7.2) and gap-only sites were removed. Tree were reconstructed using iqtree3.0^27^ (-m MFP -B 1000) and visualised using iTOL^28^.

### Ecological and statistical analyses

Analyses were performed in R (v4.4.3; R Core team). Alpha-diversity indices were computed using Microbiome^29^ (v1.28). Differences between sites were tested with a Kruskal-Wallis followed by Wilcoxon *post-hoc* test using Rstatix^30^ (v0.7.3, Bonferroni-adjusted *p*-values). Beta-diversity analyses were conducted using Jaccard and Bray-Curtis dissimilarities using Vegan^31^ (v.2.7-2). Effect of water treatment (raw *vs*. treated water), water system network (RW and TRW *vs*. GW and TGW), sampling weeks and their interactions were tested using PERMANOVA (*adonis*, Vegan^31^). Pairwise comparisons were assessed when the sampling design was balanced using pairwiseAdonis^32^ (v0.4.1). Dispersion between RW and TW was assessed by comparing the homogeneity among pairwise Jaccard dissimilarity values (Levene test, Stats, Rbase). Only pairwise comparisons within a site were considered. SIMPER analyses were performed to identify genera that explained the composition differences among sites (*simper*, Vegan^31^). Differences in relative abundances and absolute quantification were compared before and after treatment for the two water system networks separately using a Wilcoxon test. *Legionella* and *Pseudomonas* species pathogenicity was assessed by literature review^33,34^ (**Table S11**). A *p*-value <0.05 was considered significant. Mean and standard deviation are indicated if not mentioned otherwise.

## Results

### Seasonal and treatment effects shape the physicochemical profiles of river and ground water sources

The RW source was characterized by a higher average temperature compared to the GW source (16.6 ±6 .1 °C *vs.* 12.5 ±0.8 °C) and strong seasonal changes (7.8 °C in February to 24 °C in July). The total organic carbon (TOC) concentrations (2.6 ±0.4 *vs.* 0.4 ±0.04 mg.L^-1^) and water conductivity were also higher in RW as compared to GW (**Table S1**). In contrast, the NO_3_ ^-^ ion concentration was nearly two times higher in GW than in RW. Generally, in GW the physicochemical parameters showed lower variability than in RW, due to reduced exposure to external conditions and the effect of soil filtration. After treatment, the concentration of PO ³ ions increased compared to raw water for both sources (**Table S1**), to prevent the potential release of lead and copper from pipes. However, the free chlorine concentration, was approximately twice as high in TRW than in TGW (0.63 ±0.07 *vs.* 0.36 ±0.03 mg.L^-1^; **Table S1**) in agreement with the higher addition of chlorine to RW due to the longer distance the water must travel to reach the reservoir after leaving the treatment plant.

### Treatment induced a marked reduction in prokaryotic diversity in groundwater compared to river water

The GW prokaryotic community harboured a significantly higher richness, compared to RW (*p*<0.01; **Fig. 1A; Table S12**). Also, the evenness and the Shannon diversity index indicated a significant higher diversity in GW (*p*<0.01; **Fig. 1A, S3B**; **Table S12**). After GW treatment, richness decreased sharply to 136 ±55.9 ASVs (873 ASVs, 339 genera in TGW) together with Shannon diversity (*p*<0.01; **Fig. 1A, S3B**; **Table S12**) despite the higher overall diversity, whereas TRW richness stayed stable compared to RW (329 ±145 ASVs, total: 1,793 ASVs, 479 genera; **Fig. 1A; S3B**), but a significant evenness decrease and a significant increase of the Berger-Parker dominance index was observed after treatment for both sources. Thus, the prokaryotic communities in the treated water were less diverse and dominated by a few abundant genera.

**Fig. 1:**
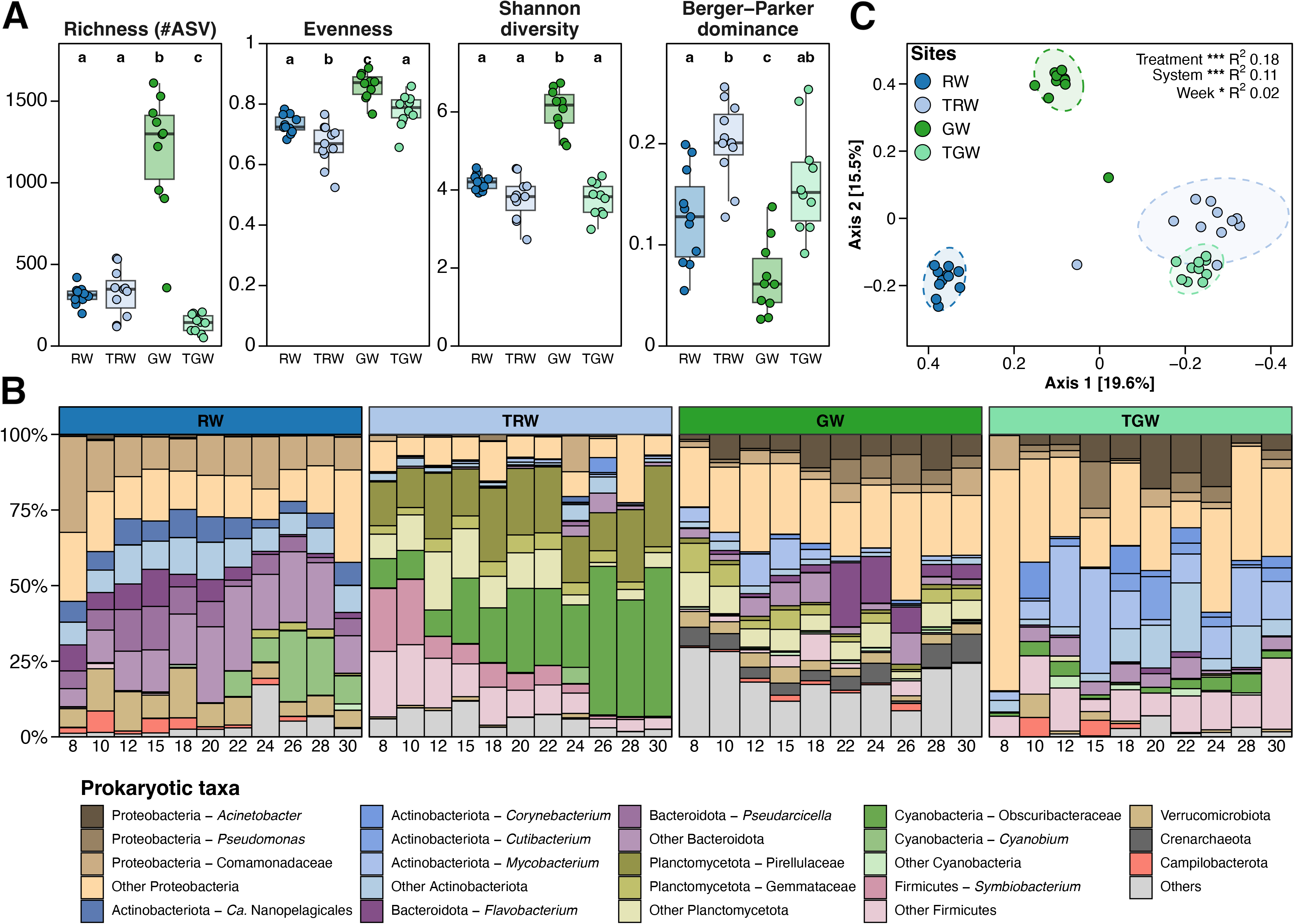
Prokaryotic community diversity and composition among water sources and their corresponding treated waters. **A**: Alpha-diversity indices (Richness, Shannon diversity index, Pielou’s evenness and Berger-Parker dominance index) based on prokaryotic ASVs. Letters refer to statistical differences using Kruskal-Wallis and Wilcoxon *post-hoc* tests (Bonferroni adjusted *p*-values). : Relative abundance (% of 16S rRNA amplicon sequencing reads) of the 10 most abundant phyla, along with most abundant genera per environment (n= 16 total, see Methods). **C**: PCoA plot (Jaccard dissimilarity) based on prokaryotic genera (see FS3 for ASV-based PCoA). Ellipses represent 95% confidence intervals and permanova statistics are displayed (treatment: raw *vs*. treated water; System: river (raw and treated) *vs*. groundwater (raw and treated); Week: 8-30). Stars refer to statistical significance (* *p*<0.05, ** *p*<0.01 and *** *p*<0.001).

### Water treatment strongly reshapes prokaryotic community composition and reduces dissimilarity among samples

In the RW and GW sources the prokaryotic communities were dominated by Proteobacteria, Bacteroidota, Actinobacteriota and Verrucomicrobiota (**Fig. 1B; Table S13**). In GW we identified in addition a higher abundance of Planctomycetota (12.6 ±5% vs. 0.9 ±1%) and Crenarchaeota (5.1 ±2% vs. 5.3×10^-2^ ±6×10^-3^%), and in the RW *Cyanobium* was identified mainly after the second half of May and showed a clear seasonal pattern (**Fig. 1B; Table S13**). However, the main composition difference between RW and GW was the higher relative abundance in RW of Comamonadaceae (*p*<0.001, 6.27% of the composition difference; **Table S13, S14**), as well as a higher relative abundance of *Cyanobium* and *Pseudarciella* (*p*<0.001, together >5% of the composition difference; **Table S13, S14**).

After RW treatment, the prokaryotic community shifted strongly and significantly due to a decrease of Comamonadaceae, an increase of Pirellulaceae, and a switch within the Cyanobacteria phylum from *Cyanobium* to Obscuribacteraceae (*p*<0.001, all >5% of the composition difference; **Fig. 1B**; **Table S13, S14**). By contrast, the change in prokaryotic communities between GW and TGW was less pronounced, with several abundant genera that persisted in GW after treatment (**Fig. S3C**). Only the higher abundance of *Mycobacterium* after treatment explained more than 5% of the composition difference (*p*<0.001; **Fig. 1B; Table S13, S14**). This suggests that the GW treatment process had a lesser effect on the prokaryotic composition (**Fig. 1B, C**). Taken together, water treatment increased the relative abundance of Firmicutes and Obscurobacteriales while decreasing Bacteroidota, regardless of the water source type.

The prokaryotic community composition of the four studied sites was significantly different, but treated samples (TRW and TGW) clustered more closely than untreated samples, suggesting that treatment leads to a partial convergence of the prokaryotic community composition, as reflected in the PCoA plot and assessed when comparing the dispersion between source and treated samples (*p*<0.01, **Fig. 1C; Table S15**). Consistently, treatment (raw *vs*. treated samples) explained a larger proportion of the variance than the water system (RW with the associated treated water vs GW with the associated treated water; *p*<0.01, R^2^ 0.18 *vs* 0.11) or temporal effects (*p*<0.05, R^2^ 0.02; **Table S15**). However, since both sources underwent different treatment processes, the effects of water source and treatment type cannot be completely disentangled.

### *Legionella*, *Pseudomonas* and *Mycobacterium* spp. persist, or even increase, after water treatment

The main waterborne opportunistic pathogens *Legionella* spp., *Pseudomonas* spp. and *Mycobacterium* spp. were detected at all studied sites, accounting for 7.0 ±9.7% of the reads (**Table S13**). Importantly, after treatment of the RW, the relative abundance of *Pseudomonas* did not decrease but stayed stable (1.91 ±3.27%), that of *Legionella* increased significantly more than 10-fold and that of *Mycobacterium* spp. showed a significant 10-fold rise in both systems (**Fig. 2A**). Considering absolute pathogen loads, the concentration of *Mycobacterium* spp. increased significantly in TRW (17.1 ±16.8 *vs*. 4.8×10^2^ ±6.2×10^2^ copie.L^-1^; **Table S16**), mirroring the rise observed in its relative abundance, whereas in GW the copy number decreased slightly but remained detectable in all TW samples (**Fig. 2B**; **Table S16**). For *Legionella*, treatment caused a significant reduction in concentration for both water sources (4.8×10^5^ ±7.5×10^5^ *vs.* 1.8×10^4^ ±4.3×10^4^). Nevertheless, *Legionella* was detectable in all TRW samples at relevant concentrations and remained detectable in TGW (**Fig. 2B**; **Table S16**). Furthermore, *L. pneumophila* and *P. aeruginosa* were detected in more than half of the water source samples and were detected at least three times after each treatment plant.

**Fig. 2:**
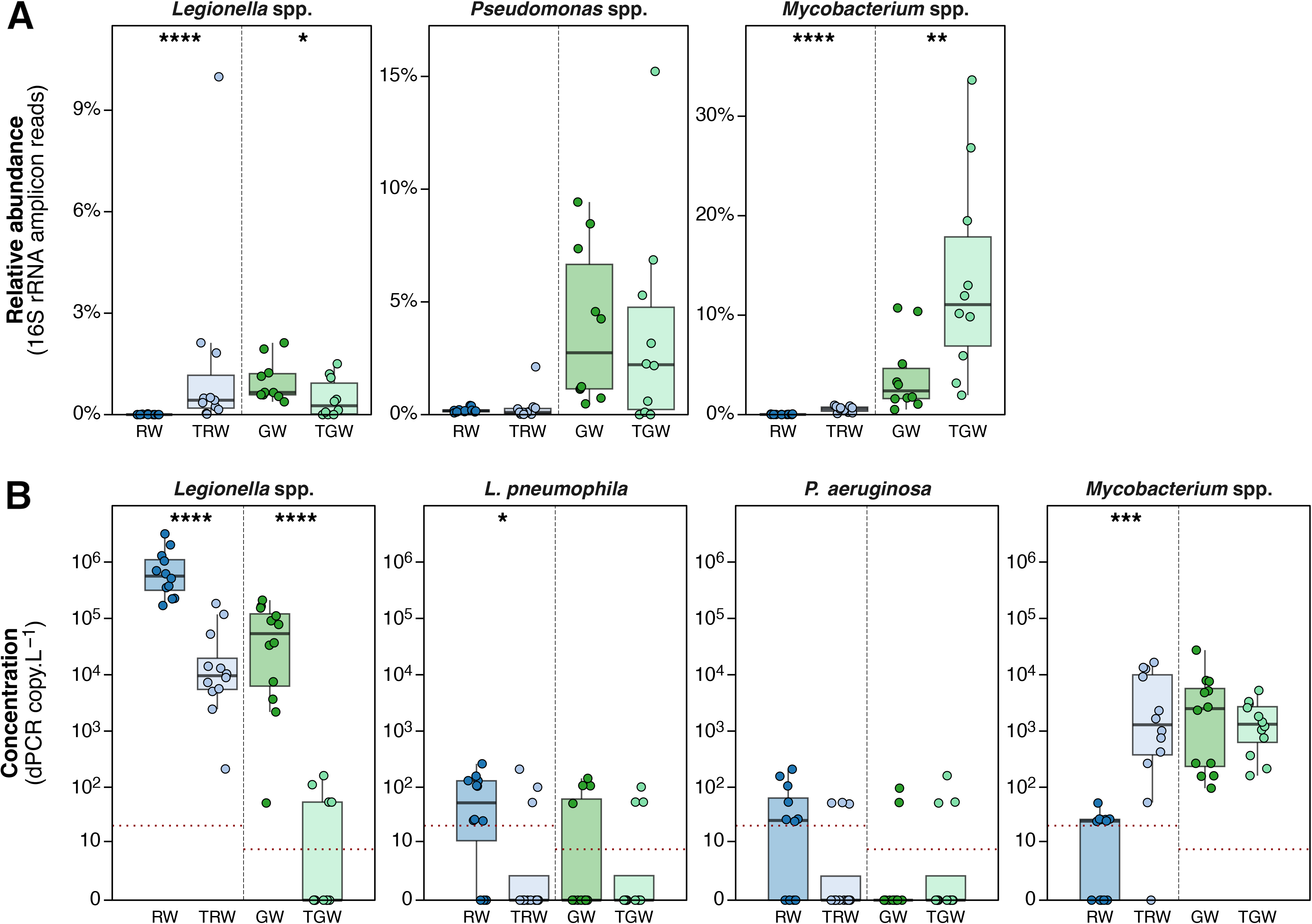
Bacterial waterborne opportunistic pathogens relative and quantitative abundances before and after water treatment processes. **A**: Relative abundance (% of 16S rRNA amplicon sequencing reads) of the *Legionella*-, *Pseudomonas*- and *Mycobacterium*-affiliated ASVs (based on SILVA 138.1). **B**: Absolute quantification (gene copy numbers per liter of filtered water) of *Legionella* spp (23S rRNA), *L. pneumophila* (*mip*), *P. aeruginosa* (*ecfX*) and *Mycobacterium* spp (*atpE*). The red line corresponds to the quantification threshold (see Methods). The y-axis is log-scaled using a log(x+1) transformation for visual purposes. Only comparisons between the same source type were computed and statistically tested (RW *vs*. TRW and GW *vs*. TGW) using Wilcoxon tests. Stars refer to statistical significance (* *p*<0.05, ** *p*<0.01, *** *p*<0.001 and **** *p*<0.0001).

### Natural and artificial aquatic environments harbour a large uncharacterized *Legionella* diversity

To obtain a deep taxonomic resolution within the genus *Legionella*, we used the *Legionella-* targeting primers 17F-28R^18^. These primers exhibited lower specificity in raw sources water than in treated water samples (38.5 ±31.3% *vs*. 71.5 ±30.7% of *Legionella* reads). After removing non*-*targeted reads, 4,625 *Legionella* ASVs (ASVs_L_ hereafter) remained. They were genetically diverse (95.3 ±1.5% seq. identity; **Fig. S1B**) and rarefaction curves reached saturation for all samples (**Fig. S4A**), indicating that our strategy captured most of the genetic diversity of *Legionella* present in the studied environments. The ASV_L_ richness was one order of magnitude higher in GW (546 ±375 per sample, 3,399 total) and RW samples (268 ±234 ASVs_L_, 1,733 total) as compared to treated water samples (TRW: 20 ±9 ASVs_L_, 114 total; TGW: 1 and 3 ASVs_L_; **Table S7**). Furthermore, 97.5% of the ASVs_L_ were found only in water sources while only 1.8% were specific to treated waters (**Fig. S4B**), indicating that water treatments drastically reduced the *Legionella* diversity.

ASVs_L_ sequences were blasted against a custom database of curated *Legionella* 16S rRNA encoding gene sequences (see Methods) to reach species-level resolution. Surprisingly, of the 4625 ASVs_L_, 67.14% (representing more than 50% of the reads in RW, TRW and GW) were distantly related to known *Legionella* species (<97% identity) but still more closely related to the genus *Legionella* or metagenome-derived Legionellaceae than to other Legionellales (*C. burnetti* and *R. massiliensis*; **Fig. 3, 4A; S1A**). Furthermore, 98.0% of those distantly related ASVs_L_ were detected in source samples. In contrast, only 1.36% ASVs_L_(15%) were very closely related to a known *Legionella* species (>99% identity; **Fig. 4A**; **Table S7**). These included *L. lytica*, *L. longbeachae*, *L.gratiana-cincinnatiensis*, *L. parisiensis*, *L. tucsonensis*, *L. londiniensis* and *L. dumoffi* as well as *L. pneumophila*, *L. rowbothamii, L. bozemanae*, *L. jeonii*, *L. maceachernii*, *L. feelei-donaldsonii* among the most abundant ASVs_L_ (**Fig. 4B; Table S7**). ASVs_L_ very closely related to pathogenic *Legionella* species were significantly more abundant in treated water than source samples (36.4 ±38.7 *vs*. 1.3 ±2.0%, *p*<0.05; **Fig. S4C**). In contrast, ASVs_L_ very closely related to non-pathogenic species were significantly less abundant in treated water samples (0.26 ±0.6% *vs*. 2.65 ±2.1%, *p*<0.05; **Fig. S4C; Table S7**). Additionally, 25.3% of the ASVs_L_ were closest to a (meta)genome-derived *Legionella* or Legionellaceae species such as DDPF01 sp021849905 among the most abundant ASVs_L_ (22.0 ±19% of the reads; **Fig. 4B, S4C**).

**Fig. 3:**
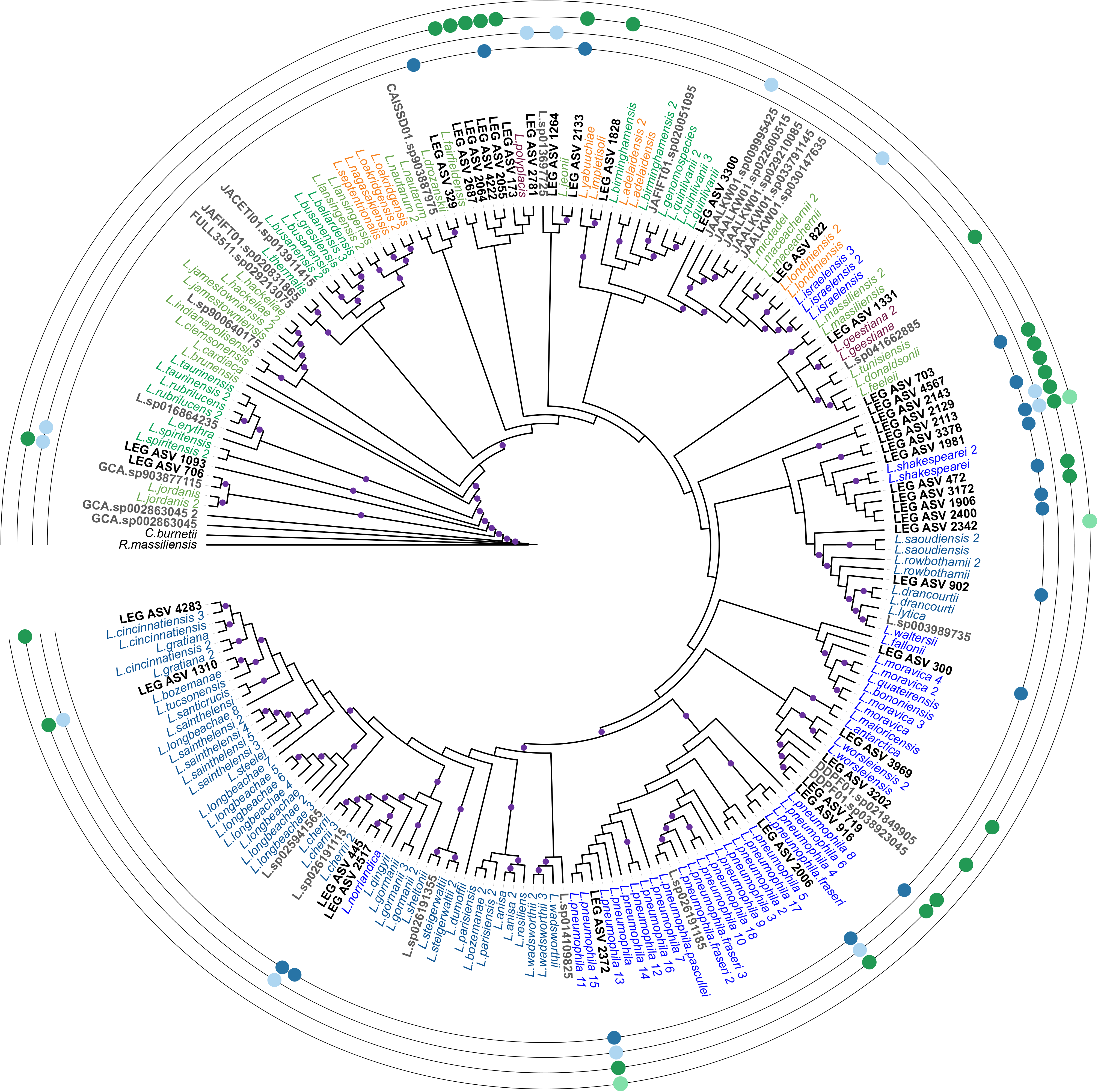
Phylogenetic relationship of the most abundant *Legionella* ASVs. Phylogenetic tree based on 17F-28R trimmed 16S rRNA encoding gene sequences from Legionellaceae from a custom database and the most abundant *Legionella*’s ASVs (ASVs_L_). Colors refer to the five clades identified from the *Legionella* core-genome phylogeny (Gomez-Valero et al. 2019). Most abundant ASVs_L_ per site are displayed in black and bold font. Colored circles on external rings indicate the presence or absence of each ASVs_L_ at each site. Bootstraps of minimum 80 are displayed.

**Fig. 4:**
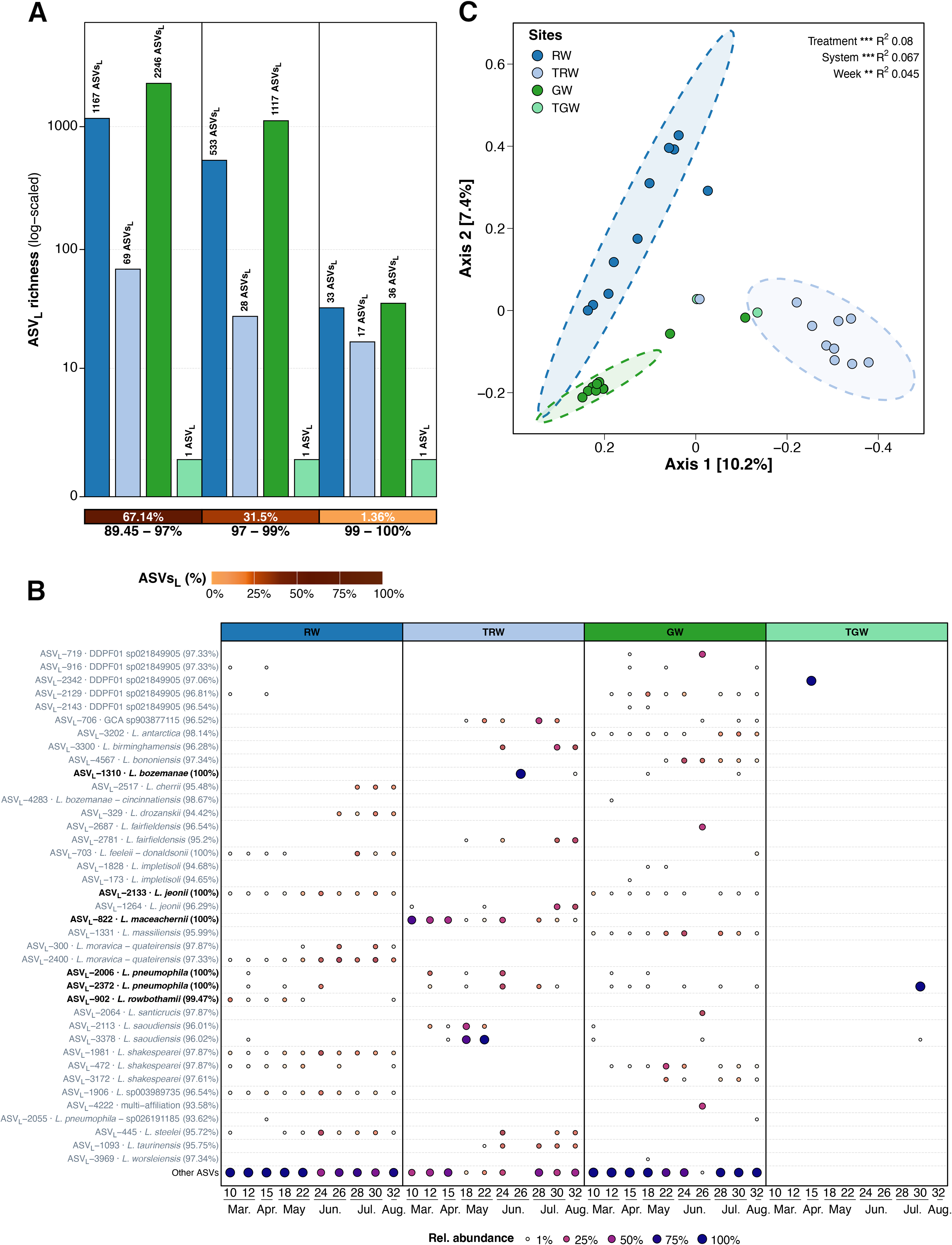
*Legionella* ASVs relatedness to known species, community diversity and composition. **A**: *Legionella* ASVs (ASVs_L_) relatedness to known *Legionella* species. The sequence identity intervals (in black) are based on intra- & inter-species sequence comparison (distantly related: <97%; closely related: 97-99%; very closely related: >99%; see Methods). The percentage of ASVs_L_ in each sequence identity interval is displayed as a heatmap and in white text. The number of *Legionella* ASVs for each at each site is indicated as a barplot (log-scaled y axis). **B**: Relative abundance of the most abundant ASVs_L_ (39 ASVs) at each site from February to July (weeks 10-32). Bold annotation refers to ASVs_L_ highly related (>99%) to only one *Legionella* species. Circles color and size refer to relative abundance. **C**: PCoA plot (Jaccard dissimilarity) based on ASVs_L_. Ellipses represent 95% confidence intervals and permanova statistics are displayed (Treatment: raw *vs*. treated water; System: river (raw and treated) *vs*. groundwater (raw and treated; Week:8-32). Stars refer to statistical significance (* *p*<0.05, ** *p*<0.01 and *** *p*<0.001).

Since most of the abundant ASVs_L_ were not closely related to any known *Legionella* species, we performed a phylogenetic analysis to determine their association with previously defined phylogenomic clades of the genus *Legionella*^35^ (**Fig. 3; Table S7**). The *L. pneumophila* clade (clade V) regrouped 26% of the most abundant ASVs_L_, whereas only 10% and 15% ASVs_L_ were related to the *L. longbeachae* (clade IV) and *L. brunensis* (clade II) clades, respectively. When considering all ASVs_L_ with at least 97% identity to an available *Legionella* sequence, the *L. longbeachae* clade was the most represented (8.24% ASVs_L_), closely followed by the *L. pneumophila* clade (7.59% ASVs_L_; **Fig. S4D**). Collectively, *Legionella*-targeting primers revealed a high and largely uncharacterized diversity of *Legionella* spp. mostly present in raw water, while enabling the high-confidence detection of known *Legionella* species. Treatment drastically reduced this diversity and selected for ASVs_L_ very closely related to clinically relevant *Legionella* species.

### *Legionella* communities are environment-specific yet *L. pneumophila* remains ubiquitous

The *Legionella* community composition appeared site-specific as almost no 95%-confidence ellipse overlap among sampling sites was observed (**Fig. 4C**) and the majority of ASVsL (87%) were detected in one environment only (**Table S7**). Also, the *Legionella* community composition of treated water samples was clearly separated from those found in natural waters as seen on the first PCoA axis (**Fig. 4C**; *p*<0.05 R^2^ 0.08). Treatment and water system explained more composition differences than temporal factors (*p*<0.005; Treatment R^2^ 0.08, system: R^2^ 0.067, week: R^2^ 0.045; **Table S15**). The most abundant ASVs_L_ were also mostly site-specific (61.5%) and many of them were only detected in natural waters (66.7%), such as the *L. rowbothamii* (ASV_L-902_) and *L. jeonii* (ASV_L-822_), never reported in disease (**Fig. 3, 4B; Table S8**). ASV_L-3202_ related to *L. antartica,* a *Legionella* species isolated from East Antarctic^36^, was primarily detected in GW samples, characterized by stable water temperatures (11.5–13.8°C throughout the sampling period), and at low abundance in river samples when the water temperature was inferior to 15°C (**Fig. 4B; Table S1, S7**).

Three ASVs_L_ most closely related to *L. shakespearei* (97.61 to 97.87% identity) were present in natural waters at most time points, but each water source contained one of the three ASVs (ASVs_L-1981_ in RW, ASVs_L-3172_ in GW) and only one was shared (ASVs_L-472_; **Fig. 4B; Table S7**). Thus, a highly different *Legionella* composition among all four studied sites was observed (**Fig. 4C, S4B**). Only two ASVs_L_ were detected at all sampling sites: one distantly related to *L. saoudiensis* (96.01% identity ASV_L-3378_) and one highly related to *L. pneumophila* (100% identity, ASV_L-2372_; **Fig. 4B**). Although this result may be biased due to the low number of ASVs from TGW, when comparing the three other sites, still only 13 of the 4625 ASVs_L_ (0.3%) were shared (**Table S7**). From the total ASVs_L_, 189 ASVs_L_ were closest to *L. pneumophila* species (95.9 ± 1.5% identity, ranging 91.8-100% and including 30 ASVs_L_ with >97% and 4 >99% identity) which were detected in almost all samples (78% of the samples, 90% in natural water sources; **Fig. S4C**, **Table S7**). These results suggest that genomic features are associated with these species that enable their adaptation to diverse environmental conditions.

Interestingly, at the whole dataset scale, the *L. pneumophila* clade was the only one detected at all sites and the most abundant one except in TRW (26.0% of the reads on average) (**Fig. S4D**). In comparison, the *L. adelaidensis* (clade I) and *L. quinlivanii* (clade III) clades each represented less than 2% of all ASVs_L_ and never exceeded 5% of the *Legionella* total reads at any site (**Fig. S4D**). The *L. brunensis* clade (clade II) cluster was intermediate, associated with 20.74% of the reads in TRW, while far less at the other sites accounting for only 4.8% of all ASVs_L_ with at least 97% identity to the database (**Fig. S4D**). Overall, ASVs_L_ related to *L. pneumophila* and other species within its phylogenomic cluster were ubiquitous.

### Water treatment shapes *Pseudomonas* communities

The *Pseudomonas*-targeting primers detected *Pseudomonas* in a similar number of samples as the universal 16S bacterial primers (464F/665R) but were highly specific (99.2 ±2.5% of the reads; **Table S9, S17**). A total of 369 ASVs (hereafter referred to as ASV_P_) were classified within the Pseudomonaceae family. These ASVs_P_ were genetically diverse with an average of 93.1 ±2.6% seq. identity among them and rarefaction curves reached saturation for all samples (**Fig. S2B, S5A**). The majority of ASVs_P_ (71%) were very closely related to known *Pseudomonas* species (>=99% seq. identity) and only 1.63% were distantly related (<95% seq. identity, see Methods; **Fig. 5A; Table S9**). Thus, based on the targeted amplicon sequences, only limited uncharacterized diversity seems to remain within this group in the habitats examined.

**Fig. 5:**
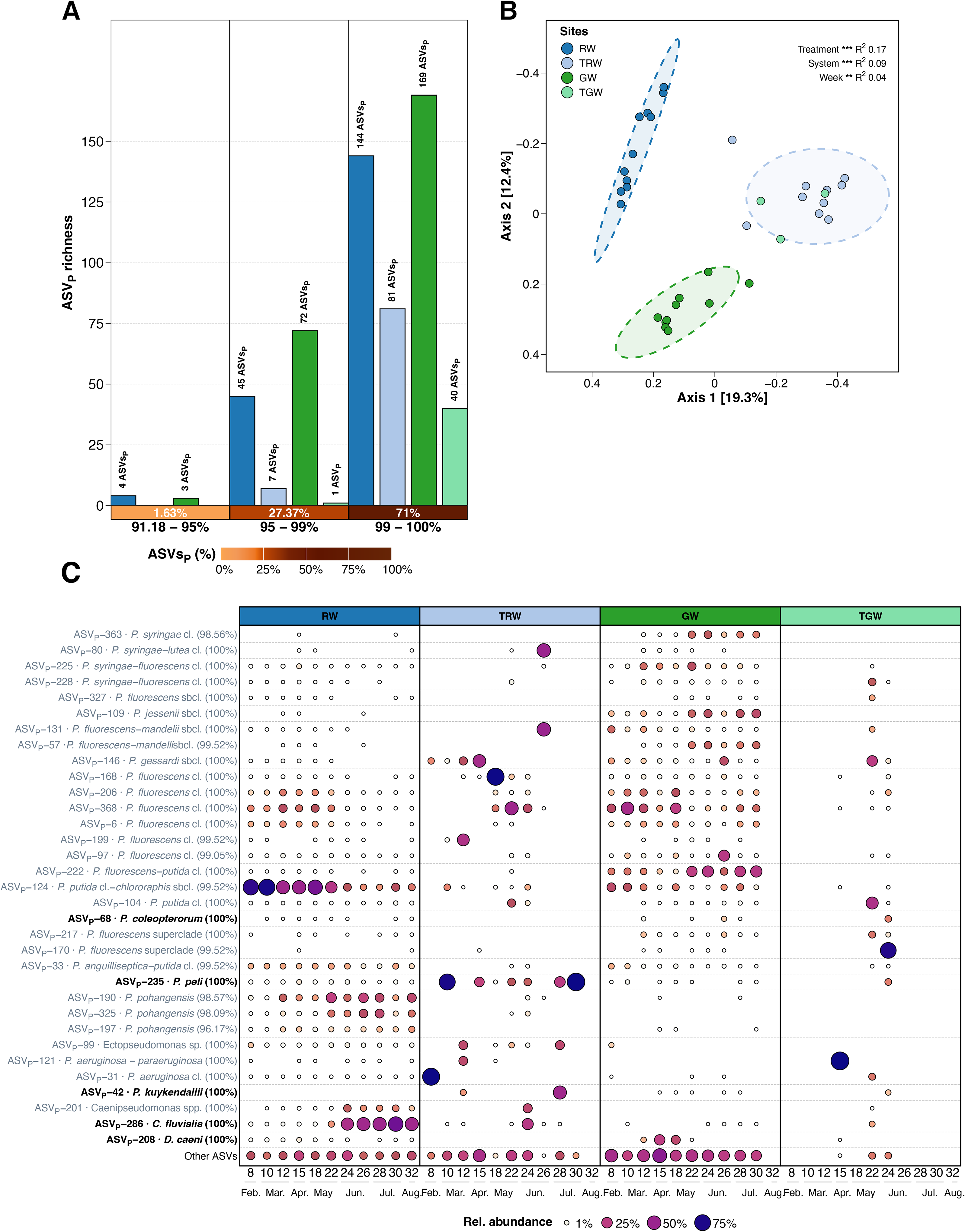
*Pseudomonas* ASVs relatedness to known species, community diversity and composition. **A**: *Pseudomonas* ASVs (ASVs_P_) relatedness to known *Pseudomonas* species based on 464F-665F trimmed 16S rRNA encoding gene seq. identity: distantly related (<97%), closely related (95-99%) and very closely related (>99%). The seq. identity intervals (in black) are based on intra- & inter-species sequence comparison (see Methods). The percentage of ASVs_P_in each seq. identity interval (in white text) is displayed as a heatmap. The number of ASVs_P_ for each interval at each site is indicated as a barplot. **B**: PCoA plot (Jaccard dissimilarity) based on ASVs_P_. Ellipses represent 95% confidence intervals and Permanova statistics are displayed (treatment: raw *vs*. treated water; Source: river (raw and treated) *vs*. groundwater (raw and treated); Week:1-12). Stars refer to statistical significance (* *p*<0.05, ** *p*<0.01, *** *p*<0.001 and **** *p*<0.0001). **C**: Relative abundance of the most abundant ASVs_P_ (33 ASVs) at each site from February to July (weeks 8-32). Bold annotation refers to ASVs_p_ highly related (>99%) to only one *Pseudomonas* species. Circles color and size refer to relative abundance.

The majority of the 369 ASVs_P_ (71.5%) was concentrated in the water sources and particularly in GW (**Fig. 5A, S5B**; **Table S9)** and most *Pseudomonas* ASVs were site-specific (67.2%), though a higher fraction was shared across all environments (5.4%, 20 ASVs_P_**; Table S9**). Among the most abundant ASVs_P,_ 39% were detected at all sites (**Table S9)**. The number of ASVs_P_ decreased after treatment in both RW and GW, as observed for *Legionella* (**Fig. 5A**; **Table S9)**. The number of ASVs that persisted after treatment was three times higher for RW than for GW, suggesting that GW treatment against *Pseudomonas* impacted diversity more (**Fig. 5A**; **Table S9)**. As displayed on the PCoA plot **(Fig. 5B**), water treatment was also the most explanatory factor of the ASVs_P_ composition (*p*<0.001, R^2^ 0.16) compared to the water system or temporal factors (*p*<0.01, R^2^ 0.09 and 0.04; **Table S15**) and as observed previously for *Legionella* it brought *Pseudomonas* communities closer together.

### Species-level resolution reveals persistence of clinically relevant *Pseudomonas* species and clades

To reach a clade- or species-level taxonomic resolution, we performed a similarity search against a custom Pseudomonadaceae database, however, the large number of described species made it difficult to assign ASVs_P_ to a single one (**Table S9**). Consequently, most ASVs_P_ were assigned to a particular clade or sub-clade. Most of the ASVs_P_ (52.3%, corresponding to 59.6 ±37% of *Pseudomonas* reads) belonged to the *P. fluorescens* superclade (**Fig. S5D; Table S9**). Within this superclade, the majority of the ASVs_P_ belonged to the *P. fluorescens* (26.6% total ASVs_P_, 30.3 ±27.6% of reads), *P. putida* (7.05% ASVs_P_, 2.3 ±7.0% of reads) and *P. syringae* (3.79% ASVs_P_, 3.4 ±9.2% of reads) clades (**Fig. S5D; Table S9)**. Among ASVs_P_ detected in all environments, 75% belonged to this superclade.

Interestingly, an association between taxonomic shifts and seasonal patterns was also observed in many of these clades. For example, in RW, the *Pseudomonas* community was dominated by *P. fluorescens* and *P. flurorescens*-*putida* clades affiliated ASVs_P_ that were dominant from February to June and then were replaced by *P. pohangensis* and *C. fluvialis* affiliated ASVs_P_ (**Fig. S5D; Table S9**). In GW, the *Pseudomonas* community was dominated by ASVs_P_ affiliated to the *P. fluorescens* clade until May, then ASVs_P_ affiliated to the *P. flurorescens*-*mandelii* and *P. flurorescens*-*jessenii* subclades and *flurorescens*-*putida* clades abundance increased (**Fig. S5D; Table S9**).

Although for most of the ASVs_P_ we could only assign them to a particular clade or sub-clade, 24% could be assigned to a single species with >99% seq. identity (**Table S9**). From these, 29%, such as ASV_P-40_ (*P. pohangensis*), were exclusively found in raw water environments while ASV_P-215_ and ASV_P-264_ (*P. fluorescens*) was detected only in GW (**Table S18**). In contrast, ASVs_p_ affiliated to three species were found exclusively in treated waters: *Halomonas pelagia*, *Pseudomonas taetrolens* and *Halomonas urumqiensis* (**Table S18**). The 90 ASVs_P_ with clear species-level assignment also include ASV_P-235_ (*P. peli*), ASV_P-286_ (*C. fluvialis*) and ASV_P-232_ (*P. japonica*) among the most abundant ASVs_P_ (**Fig. 5C; Table S9**). ASV_P-235_ was abundant in most TW samples and ASV_P-286_ was detected at all sites (**Fig. 5C; Table S9**). *P. aeruginosa*, an important human pathogen inside the *Pseudomonas* group, was not unequivocally identified, but five of the ASVs_P_ assigned to the aeruginosa clade matched to both *P. aeruginosa* and *P*. *paraeruginosa* (>99.5% seq. identity, and the most abundant one was detected at all sites except GW (ASVs_P-121_, 100% seq. identity) (**Fig. 5C, S5D; Table S9**). Although *P. aeruginosa* and *P*. *paraeruginosa* ASVs_P_ were not detected in GW samples, 3 out of 5 were detected after treatment in TGW. Moreover ASVs_P-31_ assigned to *P. aeruginosa* clade (100% seq. identity), was among the most abundant and detected at all sites (**Fig. 5C, S5D; Table S9**). All together our results suggest that *P. aeruginosa* and close species are highly adaptable to different habitats and in particular to drinking water systems.

### Water treatment decreases eukaryotic diversity and reshapes the community composition

The eukaryotic richness was statistically similar between the water sources, with 421 ±216 ASVs in RW (2,972 total ASVs, 388 genera) and 590 ±263 (1,624 total ASVs, 346 genera) in GW (*p*>0.05; **Fig. 6A; Table S4, S12**). The same trend was observed for their evenness and Shannon diversity at the ASV and genus levels (**Fig. 6A**). After water treatment, eukaryotic richness and Shannon diversity decreased significantly for samples from both sources (*p*<0.05; **Fig. 6A; Table S12**). The evenness also decreased from water to treated sources, but not significantly for the groundwater network (**Fig. 6A**). Water treatment explained more differences in the eukaryotic composition than water systems or temporal factors, as seen on the first axis of the PCoA (*p*<0.01; R^2^ 0.12, 0.08 and 0.03 respectively; **Fig. 6B; Table S15**). After treatment, the eukaryotic communities shared more taxa, thus became more similar (*p*<0.05; **Table S15**), although their most abundant taxa differed as displayed on the Bray-Curtis PCoA plot (**Fig. S6A**). For detailed community composition analysis (**Fig. 6C**) see supplementary materials.

**Fig. 6:**
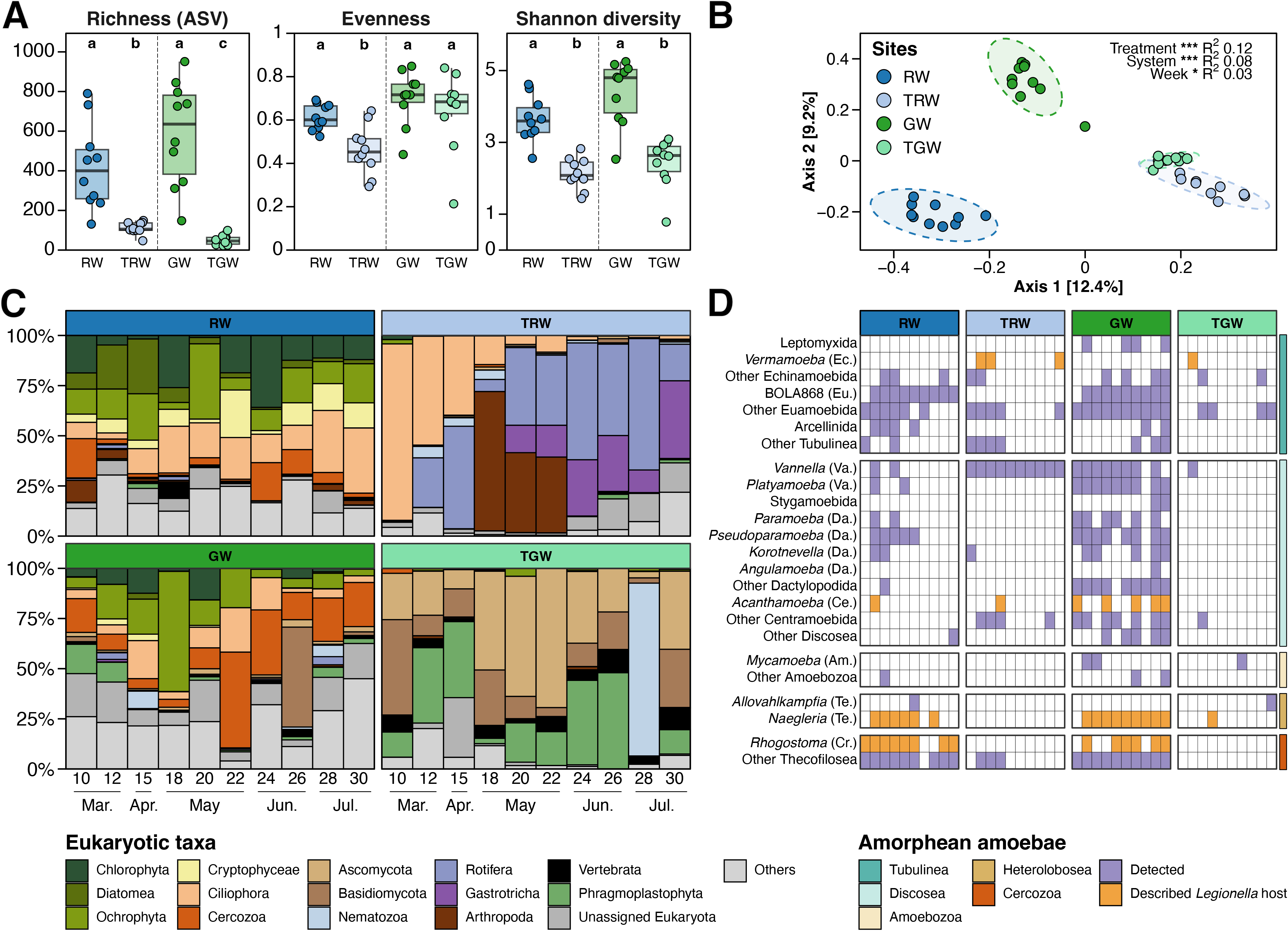
Eukaryotic community diversity and composition among water sources and their respective treated waters. **A**: Alpha-diversity indices (Richness, Pielou’s evenness and Shannon diversity index) based on eukaryotic ASVs. Letters refer to statistical differences using Kruskall-Wallis and Wilcoxon *post-hoc* tests (Bonferroni adjusted *p*-values). **B**: PCoA plot (Jaccard dissimilarity) based on prokaryotic ASVs. Ellipses represent 95% confidence intervals and permanova statistics are displayed (treatment: raw *vs*. treated water; system: river (raw and treated) *vs*. groundwater (raw and treated); week:10-30). Stars refer to statistical significance (* *p*<0.05, ** *p*<0.01 and *** *p*<0.001). **C**: Taxonomic composition (SILVA 138. phylum rank). The 10 most abundant phyla are displayed. **D**: Detection of amorphean amoebaean taxa (Solbach et al. 2021) among samples. The right-side colorbar refers to SILVA Phyla and Orders are indicated on the y-axis labels as: Echinamoebida (Ec.), Euamoebida (Eu.), Vannellida (Va.), Dactylopodida (Da.), Centramoebida (Ce.), Amoebozoa (Am.), Tetramitia (Te.) and Cryomonadida (Cr.). Described *Legionella* hosts are highlighted (Solbach et al. 2021).

Among eukaryotes, amoeba taxa accounted for 4.01 ±8.73% of the reads. At least one of the genera containing known host species for *Legionella* (*Acanthamoeba, Naegleria, Vermamoeba, Rhogostoma*) was detected in more than 50% of water source samples and at all sites (**Fig. 6D**). Interestingly, *Acanthamoeba* and *Naegleria* were only detected once in TW samples, suggesting that treatment generally removes or efficiently reduces these amoebae. In contrast, *Vermamoeba* (Tubelinea), frequently reported as one of the most abundant free-living amoebae in drinking-water systems^37^ was exclusively detected in treated water samples (**Fig. 6D**). Finally, *Vanella* (Discosea) was detected at all sites, including all TRW in and nearly all GW samples, suggesting both, high prevalence in natural environments and adaptability to artificial ones (**Fig. 6D**). Overall, water treatment processes strongly reduced the eukaryotic diversity and reshaped its communities from usual freshwater taxa to macro-eukaryotes or fungi dominated communities, while also allowing the persistence or emergence of taxa that can act as reservoirs or vectors for opportunistic pathogens.

## Discussion

Our study analysing simultaneously prokaryotic and eukaryotic communities in the river or groundwater fed drinking water systems of Paris showed that GW samples had a higher prokaryotic richness and diversity compared to RW. This is different to some previous studies that reported contrasting patterns^38–40^, likely reflecting differences in source water characteristics and analytical methodologies. Here, the analyzed GW originated from the convergence of multiple aquifers before entering the treatment plant, which may explain its higher prokaryotic richness. In contrast, despite this convergence, the eukaryotic richness and diversity of GW remained similar to those of RW samples. This is largely due to the limited or complete absence of light, which restricts primary production and favours chemosynthesis-based communities, and to carbon compound limitation that influences growth of microorganisms.

Interestingly, we show that water treatment reduced both prokaryotic and eukaryotic richness and diversity in treated water from both sources, with a greater reduction in prokaryotic diversity in GW than in RW, which differed from previous reports that did not observe an effect of water treatment on α-diversity indices^41^ or found a general decline across water source types^42^. Our results further suggest that source-specific responses may be masked in large-scale syntheses, as reported previously^42^, but might become evident in comparative studies when contrasting source and treated waters are evaluated using a common analytical framework, as presented here. The community composition in RW and GW was generally consistent with previous studies of river and groundwater systems, but here GW samples showed a higher abundance of Planctomycetota than typically reported for other aquifers. The main compositional difference between both RW and GW was the greater abundance of Comamonadaceae in RW, consistent also with observations from downstream sites of the nearby Seine River^43^.

Treatment strongly modified the community composition of both water sources. In the RW system, increasing the relative abundance of Obscuribacteraceae and Pirellulaceae, as well as the family Gemmataceae, all of which have previously been associated with sand filters^44^ used during treatment. By contrast, changes in prokaryotic community composition between GW and TGW were less pronounced, suggesting that GW treatment had a smaller effect on the prokaryotic composition. Overall, water treatment increased the relative abundance of Firmicutes and Obscurobacteriales while decreasing Bacteroidota, regardless of the water source. A treatment processes including chlorination has been reported to reduce Proteobacteria and to promote Firmicutes in treated water^45^ with Proteobacteria remaining the dominant phylum after treatment in other drinking water studies^41,46,47^. These common results highlight that treatment effects may depend on water source and treatment processes, even at the phylum level.

Our results also showed that treatment was the main driver of community shifts in both drinking water systems and for both prokaryotes and eukaryotes, although its effects depended on the water source and the nature of the treatment. While the prokaryotic communities of the two treated waters remained distinct, compositional differences were significantly reduced compared to those observed between the corresponding water sources, and a similar pattern was observed for eukaryotes. This suggests that treatment drove convergence of community composition despite differences in source water and treatment processes, likely because both treatments target the same waterborne pathogens and indirectly select for treatment-resistant taxa. As in our study we also applied a genus-specific amplicon strategy for both *Legionella* and *Pseudomonas* to reach species-level resolution we observed, consistent with our data on the bacterial composition, that treated waters shared the presence of specific opportunistic bacterial pathogens. In particular, except for *Legionella* in the GW system, the relative abundance of *Legionella*, *Mycobacterium*, and *Pseudomonas* increased or remained stable after both treatments. While *Mycobacterium* also exhibited an increase in total abundance, *Legionella*, although reduced remained detectable in all TRW samples, and *L. pneumophila* and *P. aeruginosa* persisted in a subset of samples. The persistence of these opportunistic pathogens may reflect a combination of treatment resistance, potential reintroduction, incorporation during the treatment process or regrowth in a biofilm in the vicinity of the sampling point. For example, biological activated carbon (BAC) filters used during treatment harbour established microbial communities involved in biodegradation^48^, including bacterial genera containing human pathogens^49,50^, particularly *Mycobacterium*^9^. Although downstream disinfection steps such as chlorination or ozonation are known to reduce these pathogens^9,51,52^.

Together with these bacterial opportunistic pathogens, we also detected the known and potential eukaryotic host for these bacteria. Indeed, after treatment *Vermamoeba*, the cercozoan *Rhogostoma* (Thecofilosea), recently reported as a *Legionella* host^53^ and the potentially relevant but less studied host Vanella were identified. Other eukaryotic groups that may contribute to the ecology and persistence of waterborne bacterial pathogens such as rotifers, a proposed overlooked *Legionella* vector in drinking water systems^54^ were also identified, as well as fungi that can be involved in symbiotic interactions with *Legionella*^55^ or algal taxa whose extracellular products can stimulate the growth of Gammaproteobacteria, including *Pseudomonas*^45^. These results show that accurate health risk assessment requires species-level identification as the waterborne opportunistic bacteria detected include both pathogenic and non-pathogenic species.

Both genus-targeting primers revealed sequences spanning the phylogenetic diversity of *Legionella* and *Pseudomonas*. However, whereas *Pseudomonas* ASVs were closely related to known species across all studied habitats, *Legionella* ASVs were mostly highly divergent from known species, revealing substantial uncharacterized diversity in the analyzed environments. A globally distributed and highly uncharacterized diversity has previously been reported within the order Legionellales^56^, but our study reports for the first time this high diversity specifically at the genus level in common freshwater environments. Using the same *Legionella* amplicon, Pereira *et al.* previously recovered 75 *Legionella* phylotypes based on 99% sequence similarity from treated water samples, most of which were also distantly related to known species. More recently, two studies applying the same strategy reported high uncharacterized *Legionella* diversity in polar^36^ and hypersaline^57^ environments, with 541 and 2,475 ASVs detected, respectively. However, this unknown diversity was thought to be restricted to uncommon, extreme, and poorly explored environments, but our results demonstrate that common natural and engineered aquatic environments harbour an even higher level of uncharacterized *Legionella* diversity, most of which is found in raw water, natural freshwater systems that remain understudied. Only a low proportion of detected ASVs was closely related to known *Legionella* species, but for most of them unambiguous annotation was possible revealing that species such as *L. pneumophila* and *L. jeonii* were among the most frequently detected, in agreement with a broad survey of globally distributed samples^56^. We also detected ASVs closely related to *L. antarctica*, suggesting a cold-adapted lineage and supporting the hypothesis that related sequences from freshwater, soil, and permafrost environments belong to this group^36^.

Interestingly, the most abundant species detected exclusively in raw water have not previously been associated with disease, whereas the most abundant species found in treated water include clinically relevant taxa, particularly in RW. This result suggests that resistance to treatment may be a key factor that certain species become associated with clinical cases. Overall, ASVs from treated water were less diverse but spanned most phylogenomic clusters, and were more frequently associated with the *L. brunensis* and *L. longbeachae* clusters, suggesting that the *Legionella* core-genome phylogeny may reflect ecological adaptation to specific environments. *L. pneumophila* (100% identity; ASV_L_-2372) was among the two ASVs that were detected across all sampling sites. The ubiquitous detection of ASVs related to *L. pneumophila* and other species within its phylogenomic cluster suggests a high adaptability of this group to diverse environments, which may partly explain the predominance of *L. pneumophila* in clinical cases and its association with human disease.

In contrast for *Pseudomonas* the already large number of described species made assignment beyond the clade level challenging, although species-level identification was achieved for 90 ASVs (24%). Interestingly, the only three species detected exclusively in treated water, *H. pelagia*, *P. taetrolens*, and *H. urumqiensis*, have previously been isolated from algae or plant rhizosphere soils^58–60^, suggesting that they may be associated with algae that persist through treatment processes. Among the most abundant taxa across all environments, we detected the *P. fluorescens* superclade (mainly *P. fluorescens* and *P. putida* clades), unlike in cooling tower samples^61^. At the species level, we found the species *P. peli*, which has been reported to exhibit high chlorine tolerance^62^, and *C. fluvialis*, originally isolated from the Ganges River^63^ across all environments. ASVs assigned to both *P. aeruginosa* and *P. paraeruginosa* were detected in RW and in both treated water systems. Moreover, ASVs assigned to the *P. aeruginosa* clade were among the most abundant and were detected across all studied environments. Together, these results suggest that, like *L. pneumophila*, *P. aeruginosa* and closely related species are highly adaptable to diverse habitats, particularly drinking water systems contributing to their high clinical prevalence. Thus, resistance to water treatments may represent one of the key barriers determining whether environmental species can persist and ultimately become associated with human disease.

In conclusion, species-level analyses revealed persistent waterborne opportunistic pathogens after treatment indicating a potential public health risk. Additional studies should assess cell viability and post storage treatment efficiency. Despite limited sampling, coupling universal and genus-targeting primers metabarcoding proved powerful for detecting clinically relevant bacterial species and their potential eukaryotic hosts. This integrated approach provides key insights into pathogen ecology in drinking water systems in a changing climate.

## Supporting information

Suppementaty Material

Suppementaty Tables

## Data availability statement

All amplicon sequencing raw reads were deposited into the Sequence Read Archive (SRA) database under the BioProject PRJNA1495557 (see Table S3-6 for individual sample SRA accession numbers). They are available for the reviewers and will be available on open access upon publication. Scripts are available at https://github.com/PierreFoucault/.

## Authorship contribution statement

PF, AEPC: Data curation, Formal analysis, Writing – original draft. MA: Investigation. JMM: Data curation. SW: Data curation, Investigation, Writing - Review & Editing. LM: Conceptualization, Writing - Review & Editing, Supervision, Project administration, Funding acquisition. CB: Conceptualization, Writing - Review & Editing, Supervision, Project administration, Funding acquisition. LGV: Conceptualization, Data curation, Formal analysis, Writing – original draft, Supervision, Project administration, Funding acquisition.

## Acknowledgment

We thank the Eau de Paris sampling team who facilitated the sampling of the facilities. We gratefully acknowledge Sandra Manco (Eau de Paris) for technical assistance and guidance with the laboratory work. We thank the treatment plant directors for their access approval. We thank the iGenSeq sequencing platform (Paris Brain Institute - ICM, Hôpital de la, CNRS UMR 7225 – Inserm U 1127 – Sorbonne Université UM75).

