## Supplementary material for "Drinking water treatment drastically modifies multi-kingdom water communities, but clinically relevant opportunistic bacterial pathogens persist": Suppementaty Tables

4

5 Pierre Foucault^1^*, Ana Elena Pérez-Cobas^1$^*, Madalina Ababii^1§^, Jesus Marin-Miret^1^,
6 Sebastien Wurtzer^2^, Laurent Moulin^2#^, Carmen Buchrieser^1#^ and Laura Gomez-Valero^1#^ 7

8

#### 9 - Extended analyses: eukaryotic community compositon changes

10 **- Extended materials and methods**

11

12 **- Supplementary Figures**
13 **Fig. S1: Relatedness among sequences from the *Legionella* curated database and among**
14 **this study’s ASVsL.**
15 **Fig. S2:** Relatedness among sequences from the Pseudomonadaceae curated database and
16 among this study’s ASVsP.
17 **Fig. S3:** Additional prokaryotic community diversity and composition among water sources
18 and their respective treated waters

#### 19 Fig. S4: Additional *Legionella* community diversity and composition among water sources

20 **and their respective treated waters.**
21 **Fig. S5:** Additional *Pseudomonas* community diversity and composition among water sources
22 and their respective treated waters
23 **Fig. S6:** Additional eukaryotic diversity and community composition 24

#### 25 - Supplementary tables

26 Table S1: Measured physic-chemical parameters
27 Table S2: dPCR probe sequences
28 Table S3: Prokaryotic universal amplicon (515Fb-806Rb) pre-process summary
29 Table S4: Eukaryotic universal amplicon (1427F-1616R) pre-process summary
30 Table S5: *Legionella* targeted amplicon (lgsp 17F-28R) pre-process summary
31 Table S6: *Pseudomonas* targeted amplicon (Pse 464F-665R) pre-process summary
32 Table S7: *Legionella* ASVs annotation summary
33 Table S8: Most abundant *Legionella* ASVs annotation summary
34 Table S9: *Pseudomonas* ASVs annotation summary
35 Table S10: Pseudomonaceae representative species table
36 Table S11: List of pathogenic Legionellaceae and Pseudomonaceae species
37 Table S11: Prokaryotic community alpha-diversity summary
38 Table S12: Alpha-diversity analyses summary
39 Table S13: Prokaryotic community composition table
40 Table S14: Prokaryotic community SIMPER analysis summary
41 Table S15: PERMANOVA analyses summary
42 Table S16: Waterborne opportunistic pathogens dPCR analysis summary
43 Table S17: PCR- and dPCR-positive samples summary
44 Table S18: *Pseudomonas* ASVs with uniquespecies annotation
45 Table S19: Eukaryotic community composition table
46 Table S20: Eukaryotic community SIMPER analysis summary 47

#### 48 Extended analyses: eukaryotic community composition changes

49 The eukaryotic richness was statistically similar between the water sources, with 421 ±216
50 ASVs in RW (2,972 total ASVs, 388 genera) and 590 ±263 (1,624 total ASVs, 346 genera) in
51 GW (*p*>0.05; **Fig. 6A; Table S4, S12**). The same trend was observed for their evenness and
52 Shannon diversity at the ASV and genus levels (**Fig. 6A**). After water treatment, eukaryotic
53 richness and Shannon diversity decreased significantly for samples from both sources (*p*<0.05;
54 **Fig. 6A; Table S12**). The evenness also decreased from water to treated sources, but not
55 significantly for the groundwater network (**Fig. 6A**). Water treatment explained more
56 differences in the eukaryotic composition than water systems or temporal factors, as seen on
57 the first axis of the PCoA (*p*<0.01; R^2^ 0.12, 0.08 and 0.03 respectively; **Fig. 6B; Table S15**).
58 RW communities were dominated by different microalgae (Ochrophyta, 15.6 ±9.7%
59 Chlorophyta, 14.2 ±11.0%; Cryptophyceae, 8.8 ±6.7%; Diatomea, 7.9 ±9.2%) as well as by
60 Ciliates (17.5 ±9.5%) whose relative abundance (17%) increased starting from April, as
61 observed from other river ecosystems^1^ (**Fig. 6C; Table S19)**. GW contained the same most
62 abundant groups, with the higher relative abundance of Cercozoa (18.2 ±13.6%, mostly
63 Thecofilosea) and Ciliophora (9.0 ±7.4%, mostly Choreotrichia) explaining most of the
64 difference between sources (**Fig. 6C;** *p*<0.05; **Table S19, S20**). After treatment, the eukaryotic
65 communities shared more taxa, thus became more similar (*p*<0.05; **Table S15**), although their
66 most abundant taxa differed as displayed on the Bray-Curtis PCoA plot (**Fig. S6A**). For detailed
67 community composition changes analysis (**Fig. 6C**) see supplementary materials. The changes
68 from RW to TRW were explained by the presence of Rotifera (34.3 ±21.9%, mostly
69 Monogononta), Arthorphoda (14.9 ±25.0 mostly Diplostraca) and Gastrotoicha (13.56 ±14.1%,
70 mostly Chaetonotida) becoming the most dominant groups (**Fig. 6C)**. Furthermore, the relative
71 abundance of Ciliophora decreased (21.3 ±29.8%) and shifted from Choreotrichia to *Aspidisca*,
72 both within the Spirotrichea order (*p*<0.05; **Fig. 6C; Table S19, S20**). In contrast, the
73 emergence of plants (23.0 ±17.1%, Magnoliophyta) as well as the increased abundance of
74 Ascomycota (33.3+21.5%, Debaryomycetaceae and Dothideomycetes) and Basidiomycota
75 (17.7 ±13.6, Malasseziomycetes) explained the composition changes from GW to TGW
76 (*p*<0.05; **Table S19, S20**). Higher relative abundance of macro-eukaryotes after water treatment
77 in cooling towers^2^ has been previously reported and could be partially due to their
78 multicellularity.

79

#### 80 Extended materials and methods

81 **Water sample collection.**
82 Samples were collected bi-weekly from February to August 2023 at the entry (raw water) and
83 exit (treated water) of two water treatment plants alimenting the Paris drinking water. The
84 Joinville water treatment plant takes water from the Marne River (river water, RW), and
85 releases produced water (treated river water, TRW) after removal of large particles, pre-
86 ozonation, coagulation-flocculation, biological sand filtration, ozone treatment, granulated
87 activated carbon, UV treatment, and chlorination. The Haÿ-les-Roses water treatment plant
88 takes water from various groundwater sources transported through the Vanne aqueduct
89 (groundwater, GW) and releases treated water (treated groundwater, TGW) after four steps:
90 coagulation with FeCl^3+^ polymer, powdered activated carbon filtration, ultrafiltration, and
91 chlorination. Both plants add orthophosphoric acid to limit the dissolubility of heavy metal in
92 old private buildings. Temperature, total organic carbon (COT), nitrates (NO₃⁻ ions), nitrites
93 (NO2⁻ ions), orthophosphates (PO₄³⁻ ions), pH, free chlorine (free Cl₂ ions), iron (Fe^2+^ ions) and
94 water conductivity were also measured (**Table S1**).

95

#### 96 Water sample processing.

97 Water samples were collected in one-litre bottles, with addition of 20 mg of sodium thiosulfate
98 to neutralize residual chlorine in the treated water samples (TRW and TGW). The bottles were
99 stored at 4 °C and transported to the laboratory for analysis within a maximum of 12 hours.
100 Once in the lab, samples were filtered using a filtration manifold equipped with sterile
101 polycarbonate membrane filters with a pore size of 0.4 μm (Millipore®): 1L of raw water and
102 3L of produced water Filters were subsequently transferred into 2 mL tubes suitable for
103 downstream nucleic acid extraction. A filtration blank was included in each batch to monitor
104 and prevent cross-contamination during sample processing. 105

#### 106 Nucleic acid extraction.

107 A combined chemical and mechanical lysis approach was used. First, 800 µL of Trizol reagent
108 was added to induce chemical lysis, followed by mechanical disruption using a bead mill
109 homogenizer (Fisherbrand™ Bead Mill 24) for 10 minutes at a speed of 3.1 m/s at room
110 temperature. The tubes were then centrifuged, and the supernatant was collected. Nucleic acid
111 purification was carried out using a QIAsymphony automated extractor (QIAGEN) with the
112 DSP Virus/Pathogen Kit (#937055), following the manufacturer’s instructions. The extracted
113 nucleic acids were eluted in a final volume of 50 µL of elution buffer and stored at 4 °C. 114

#### 115 Absolute quantification of opportunistic waterborne pathogens.

116 Digital PCR (dPCR) were performed using the QIAcuity instrument (QIAGEN) with the
117 QIAcuity PCR Probe Master Mix Kit (#250098) on the Nanoplate 26k 24-well plate (#250102)
118 following the manufacturer’s instructions. Target-specific primers, probes and reaction mixes
119 used are listed in Table S2. The dPCR cycling protocol consisted of an initial polymerase
120 activation step at 95 °C for 2 minutes, followed by 45 amplification cycles of 95 °C for 15
121 seconds and 56 °C for 1 minute. Results were analyzed using QIAcuity Software Suite
122 v.2.1.7.182. Considering the sample concentration factor and the quantification method, the
123 limit of detection/quantification was estimated at approximately 20 copie. L^-1^ for source water
124 samples and 7 copie. L^-1^ for treated water samples. 125

#### 126 Amplicon PCR, Illumina MiSeq library preparation and sequencing.

127 Two universal primers set were used for prokaryotes (515Fb/806Rb, V4 region of the 16S
128 rRNA encoding gene)^3,4^ and eukaryotes (1427F/1616R, V8 region of the 18S rRNA encoding
129 gene^5^. Additionally, two genus-targeting primers set were used for *Legionella* (Lgsp17F/28R)^6^
130 and Pseudomonas (Pse464F/665R)^7^, both targeting the V3–V4 regions of the 16S rRNA
131 encoding gene. All primer sets contained Illumina indexing primers. PCR amplifications were
132 performed in 50 µL reaction mixtures containing 5 µL of crude DNA extract, 10 µL of Q5
133 buffer (New England Biolabs), 3% DMSO, each dNTP at a final concentration of 0.25 mM, 1
134 U of Q5 High Fidelity DNA polymerase (New England Biolabs), and 0.5 µM of each forward
135 and reverse primer. PCRs were carried out for 35 cycles, with annealing temperatures adjusted
136 for each primer set (515Fb/806Rb: 57 - °C; F1427/R1616: 64 °C; Lgsp17F/Lgsp28R: 67 °C;
137 Pse34F/Pse665R: 62 °C). Amplicons were visualized by electrophoresis on 1.5% agarose gels.
138 Sequencing libraries were prepared following the manufacturer’s instructions using the Nextera
139 XT DNA Library Preparation Kit (Illumina, #FC-131-102). Sequencing (Illumina MiSeq,
140 2x150 bp) was performed at the Institut du Cerveau (ICM) iGenSeq sequencing platform (Paris,
141 France).

142

#### 143 Amplicon sequence pre-processing.

144 Sequence pre-processing was performed using the QIIME2^8^ workflow (v2024.5).	Reads
145 without primers were discarded and trimming was performed for both forward and reverse reads
146 (515Fb/806Rb: f-203, r-108; 17F/28R: f-240, r-162; 464F/665R: f-190, r-136; 1427F/1616R:
147 f-130, r-100). Amplicon sequence variants (ASVs) were generated after quality filtering,
148 merging and chimeras’ removal with the DADA2 plugin^9^ (v1.30.0, default: error rate: 2 and
149 min. overlap: 12). A second chimera filtering step was performed using the VSEARCH^10^ plugin
150 and resulting ASVs with less than 10 counts across all samples were discarded. Taxonomic
151 affiliation was performed using the *sklearn* algorithm trained on the SILVA-138.2 database^11^.
152 Eukaryota-, mitochondria- and chloroplast-affiliated ASVs were discarded from the
153 prokaryotic,	*Legionella*	and	*Pseudomonas*	datasets*.*	Prokaryota-,	mitochondria-	and
154 chloroplast-affiliated ASVs were discarded from the eukaryotic dataset. Rarefaction was
155 performed at 4,600 and 60,000 reads for prokaryotic and eukaryotic amplicons respectively,
156 yielding 9,431 total ASVs in 42 samples for the prokaryotic dataset and 4,590 total ASVs in 40
157 samples for the eukaryotic dataset (**Table S3, S4**). For the genus-targeting datasets, sequences
158 belonging to the *Legionella* (4,625 total ASVs, dataset hereafter referred to as ASVL, 32 final
159 samples) and *Pseudomonas* (335 total ASVs, dataset hereafter referred to as ASVP, 34 final
160 samples;) genera were respectively filtered (**Table S5, S6**). For the *Legionella* dataset, samples
161 Legio-S1-rivB-S39, Legio-S1-rivB-S40, Legio-S7-souT-S70 and Legio-S9-souT-S77 were
162 discarded from all analyses including pre-process, due to the absence of *Legionella* reads.

163

164 **Curated taxonomic annotation of *Legionella* ASVs.**
165 A curated in-house strain-level database of the Legionellaceae family was created, including
166 metagenome-derived, *Tatlotkia* and former *Fluoribacter* species. 16S rRNA sequences were
167 downloaded from four existing curated databases: LPSN^12^; April 30^th^ 2025), NCBI Bacterial
168 16S Ribosomal RNA RefSeq Targeted Loci Project (October 30th 2025), MIMt 16S-M2c-24-
169 10^13^, and GTDB-r226^14^ bac120-all-ssu (including 480 non-*pneumophila* and including 67
170 metagenome-derived sequences assigned as *Legionella* or *Legionellaceae*). The 16S rRNA
171 sequence of *L. genomospecies* strain 2055-AUS-E from NCBI (txid1093625) was manually
172 added. Representative 16S rRNA sequences of *Coxellia burnetti* and *Diplorickettsia*
173 *massiliensis* from GTDB-r226^14^ bac120 ssu reps were added as outgroups from the
174 Legionellales order. Sequences affiliated to *L. fraseri* and *L. pasculeii* strains were annotated
175 as *L. pneumophila* subspecies and all species and strain names were manually checked and
176 corrected if necessary for consistency across databases. Using a custom QIIME2 script,
177 sequences were trimmed according to the *Legionella*-targeting primers Lgsp 17F-18R^6^ filtered
178 for the presence of ambiguous bases (*e.g.* N) or homopolymers longer than 8 nucleotides. The
179 remaining sequences were dereplicated at the strain level (*i.e.* only identical sequences from the
180 same species and strain names were dereplicated), resulting in 1235 strain-level unique
181 sequences (including 2 Legionellales outgroup sequences). All ASVL sequences were blasted
182 against the *Legionella* database using blastn^15^ (v2.15.0, evalue <= 1e-10). Multiple species
183 annotations were reported if the best hits had equal BIT scores (**Table S7, S8**). 184
185 **Curated taxonomic annotation of *Pseudomonas* ASVs.**
186 A curated database was created to represent the Pseudomonaceae family (including species
187 from previous *Pseudomonas* clades now described as new genera^16–20^: *Halospeudomonas* (prev.
188 *P. pertucinogena* clade)*, Stutzerimonas* (prev. *P. stutzeri* clade), *Aquipseudomonas* (prev.
189 *P. alcaligenes* clade), *Caenipseudomonas* (prev. *P. fluvialis* clade), *Ectopseudomonas* (prev.
190 *P. oleovorans* clade), *Geospeudomonas* (prev. *P. linyingensis* clade), *Metapseudomonas* (prev.
191 *P. resinovorans* clade), *Phytopseudomonas* (prev. *P. straminea* clade), *Zestomonas* (prev.
192 *P. thermotolerans* clade), *Chryseomonas* (prev. *P. oryzihabitans/P. luteola* clade) and *Serpens*
193 (prev. *P. flexibilis/alcaligenes* clade). The database was based on existing curated species-level
194 databases: LPSN^12^ (May 19^th^ 2025); GTDB-r226^14^ bac120-ssu-reps; NCBI Bacterial 16S
195 Ribosomal RNA RefSeq Targeted Loci Project (May 7^th^ 2025); MIMt 16S M2c-24-10^13^.
196 Representatives 16S rRNA sequences of *Escherichia coli* and *Cellvibrio japonicus* from
197 GTDB-r226^14^ bac120-ssu-reps were added as outgroups. Sequences were trimmed according
198 to *Pseudomonas*-targeting primers Pse 464F/665R^7^, filtered for the presence of ambiguous
199 bases (*e.g.* N) or homopolymers longer than 8 nucleotides and dereplicated at the species level
200 using a custom QIIME2 script. The final database contained 734 sequences (including 3
201 outgroup sequences). All ASVsP sequences were blasted against the database using blastn^15^
202 (evalue <= 1e-10). Multiple annotations were reported if the best hits had equal BIT scores and
203 belonged to different genome-defined genus, superclade, clade, subclade or species (**Table S9**).

204

#### 205 Intra- and inter-species threshold using 17F-28R and 4364F-665R primers.

206 No previous study established a threshold for species-delimitation for the two genus-targeting
207 primers used, thus all intra- and inter- species pairwise comparisons were computed using this
208 study’s custom Legionellaceae and Pseudomonaceae databases. For the *Legionella*-targeting
209 primers, the >= 99% sequence identity threshold was selected as representing two sequences
210 belonging to the same known species and the lowest probability to correspond to different
211 species (“very closely related”; 93.9% of intra-species comparisons *vs.* 0.5% of inter-species
212 comparisons; **Fig. S1**). The 97-99% identity interval corresponded to possibly related to a
213 known species (“closely related”; 5.6% of intra-species comparisons *vs.* 31% of inter-species
214 comparisons) while the sequences with less than 97% identity indicated sequences “distantly
215 related” to a known species (**Fig. S1**). However, for the *Pseudomonas*-targeting primers, the
216 difference between the intra- and inter-species sequence identity ranges was less clear-cut with
217 yet most intra-species comparisons were within the 95-99% sequence identity interval (**Fig.**
218 **S2**). The >=99% sequence identity threshold was thus considered as representing for sequences
219 “very closely related” to a known species (3.7% of inter-species comparisons), the 95-99 for
220 sequences “closely related” sequences and a sequence identity below 95% was considered as
221 “distantly related” to a known *Pseudomonas* species (**Fig. S2**). 222

#### 223 Sequence alignment and tree visualization of the most abundant ASVs.

224 For both *Legionella* and *Pseudomonas* datasets, the most abundant ASVs were defined as a
225 subset of the 10 ASVs with the highest relative abundance average over at each by sites (RW,
226 TRW, GW, TGW) grouped together (38 and 31 ASVs total respectively). Their sequences,
227 together with their respective species-level dereplicated sequences from curated databases,
228 were aligned using the SILVA-based aligner SINA (v1.7.2, SILVA 138.2^21^ and gap-only sites
229 were removed. A subset of the Pseudomonaceae database (162 sequences) including only
230 genome-based genus, clade and subclade representative species listed in **Table S10** was used
231 for visualisation purposes^19,20^. Tree reconstructions were computed using iqtree3.0^22^ with the
232 best-fit model according to the BIC criteria (-m MFP) and 1000 ultrafast bootstraps (-B 1000).
233 Newick files were visualised and annotated using the iTOL web browser^23^. 234

#### 235 Ecological and statistical analyses

236 Analysis, metrics computation, plots and statistical testing were performed using R (v4.4.3; R
237 Core team). Alpha-diversity indices (richness, Shannon diversity index, Pieliou’s evenness and
238 Berger-Parker dominance index were computed using Microbiome^24^ (v1.28). Differences
239 between sites were tested with a Kruskal-Wallis followed by Wilcoxon post-hoc test and p-
240 values were adjusted (Bonferroni) using Rstatix^25^ (v0.7.3). Beta-diversity analyses were
241 conducted using Jaccard and Bray-Curtis dissimilarities using Vegan^26^ (v.2.7-2). Effect of
242 water treatment (raw vs. treated water), water system network (RW and TRW vs. GW and
243 TGW), sampling weeks and their interactions were tested using PERMANOVA tests (*adonis*,
244 Vegan). Pairwise comparisons were assessed when the sampling design was balanced using
245 pairwiseAdonis^27^ (v0.4.1). Dispersion between raw and treated water samples was assessed by
246 comparing the homogeneity among pairwise Jaccard dissimilarity values using a Levene test
247 (Stats, Rbase). Only pairwise comparisons within a site were considered. SIMPER analyses
248 were performed to identify genera that explained the composition differences among sites
249 visualized in their respective PCoAs (*simper*, Vegan^26^). Differences in relative abundances
250 (*Legionella* spp., *Pseudomonas* spp. and *Mycobacterium* spp or between non-pathogenic and
251 pathogenic species) and absolute quantification (*Legionella* spp., *L. pneumophila*, *P.*
252 *aeruginosa* and *Mycobacterium* spp.) were compared before and after treatment for river and
253 groundwater samples separately using a Wilcoxon rank test. *Legionella* and *Pseudomonas*
254 species pathogenicity (either established or putative) was assessed by literature review^28,29^
255 (**Table S11**). A *p*-value <0.05 was considered significant. Mean and standard deviation are
256 indicated if not mentioned otherwise.

#### 257 References

| 258 |  | |
| --- | --- | --- |
| 259 | (1) | Potvin, M.; Rautio, M.; Lovejoy, C. Freshwater Microbial Eukaryotic Core Communities, |
| 260 |  | Open-Water and Under-Ice Specialists in Southern Victoria Island Lakes (Ekaluktutiak, |
| 261 |  | NU, Canada). *Front. Microbiol.* **2022**, *12*. <https://doi.org/10.3389/fmicb.2021.786094>. |
| 262 | (2) | Paranjape, K.; Bédard, É.; Shetty, D.; Hu, M.; Choon, F. C. P.; Prévost, M.; Faucher, S. |
| 263 |  | P. Unravelling the Importance of the Eukaryotic and Bacterial Communities and Their |
| 264 |  | Relationship with Legionella Spp. Ecology in Cooling Towers: A Complex Network. |
| 265 |  | *Microbiome* **2020**, *8* (1), 157. <https://doi.org/10.1186/s40168-020-00926-6>. |
| 266 | (3) | Apprill, A.; McNally, S.; Parsons, R.; Weber, L. Minor Revision to V4 Region SSU rRNA |
| 267 |  | 806R Gene Primer Greatly Increases Detection of SAR11 Bacterioplankton. *Aquat.* |
| 268 |  | *Microb. Ecol.* **2015**, *75*. <https://doi.org/10.3354/ame01753>. |
| 269 | (4) | Parada, A. E.; Needham, D. M.; Fuhrman, J. A. Every Base Matters: Assessing Small |
| 270 |  | Subunit rRNA Primers for Marine Microbiomes with Mock Communities, Time Series |
| 271 |  | and Global Field Samples. *Environ. Microbiol.* **2016**, *18* (5), 1403–1414. |
| 272 |  | <https://doi.org/10.1111/1462-2920.13023>. |
| 273 | (5) | Hannen, E. J. V.; Agterveld, M. P. V.; Gons, H. J.; Laanbroek, H. J. REVEALING |
| 274 |  | GENETIC DIVERSITY OF EUKARYOTIC MICROORGANISMS IN AQUATIC |
| 275 |  | ENVIRONMENTS BY DENATURING GRADIENT GEL ELECTROPHORESIS 1 The |
| 276 |  | Application of Molecular Biological Tech- Niques to Solve Problems in Microbiology |
| 277 |  | Has Changed Our View of the Diversity And. **1998**, *213*, 206–213. |
| 278 | (6) | Kahlisch, L.; Henne, K.; Draheim, J.; Brettar, I.; Höfle, M. G. High-Resolution In Situ |
| 279 |  | Genotyping of Legionella Pneumophila Populations in Drinking Water by Multiple-Locus |
| 280 |  | Variable-Number Tandem-Repeat Analysis Using Environmental DNA. *Appl. Environ.* |
| 281 |  | *Microbiol.* **2010**, *76* (18), 6186–6195. <https://doi.org/10.1128/AEM.00416-10>. |
| 282 | (7) | Bergmark, L.; Poulsen, P. H. B.; Al-Soud, W. A.; Norman, A.; Hansen, L. H.; Sørensen, |
| 283 |  | S. J. Assessment of the Specificity of Burkholderia and Pseudomonas qPCR Assays for |
| 284 |  | Detection of These Genera in Soil Using 454 Pyrosequencing. *FEMS Microbiol. Lett.* |
| 285 |  | **2012**, *333* (1), 77–84. <https://doi.org/10.1111/j.1574-6968.2012.02601.x>. |
| 286 | (8) | Bolyen, E.; Rideout, J. R.; Dillon, M. R.; Bokulich, N. A.; Abnet, C. C.; Al-Ghalith, G. |
| 287 |  | A.; Alexander, H.; Alm, E. J.; Arumugam, M.; Asnicar, F.; Bai, Y.; Bisanz, J. E.; Bittinger, |
| 288 |  | K.; Brejnrod, A.; Brislawn, C. J.; Brown, C. T.; Callahan, B. J.; Caraballo-Rodríguez, A. |
| 289 |  | M.; Chase, J.; Cope, E. K.; Da Silva, R.; Diener, C.; Dorrestein, P. C.; Douglas, G. M.; |
| 290 |  | Durall, D. M.; Duvallet, C.; Edwardson, C. F.; Ernst, M.; Estaki, M.; Fouquier, J.; |
| 291 |  | Gauglitz, J. M.; Gibbons, S. M.; Gibson, D. L.; Gonzalez, A.; Gorlick, K.; Guo, J.; |
| 292 |  | Hillmann, B.; Holmes, S.; Holste, H.; Huttenhower, C.; Huttley, G. A.; Janssen, S.; |
| 293 |  | Jarmusch, A. K.; Jiang, L.; Kaehler, B. D.; Kang, K. B.; Keefe, C. R.; Keim, P.; Kelley, |
| 294 |  | S. T.; Knights, D.; Koester, I.; Kosciolek, T.; Kreps, J.; Langille, M. G. I.; Lee, J.; Ley, |
| 295 |  | R.; Liu, Y.-X.; Loftfield, E.; Lozupone, C.; Maher, M.; Marotz, C.; Martin, B. D.; |
| 296 |  | McDonald, D.; McIver, L. J.; Melnik, A. V.; Metcalf, J. L.; Morgan, S. C.; Morton, J. T.; |
| 297 |  | Naimey, A. T.; Navas-Molina, J. A.; Nothias, L. F.; Orchanian, S. B.; Pearson, T.; Peoples, |
| 298 |  | S. L.; Petras, D.; Preuss, M. L.; Pruesse, E.; Rasmussen, L. B.; Rivers, A.; Robeson, M. |
| 299 |  | S.; Rosenthal, P.; Segata, N.; Shaffer, M.; Shiffer, A.; Sinha, R.; Song, S. J.; Spear, J. R.; |
| 300 |  | Swafford, A. D.; Thompson, L. R.; Torres, P. J.; Trinh, P.; Tripathi, A.; Turnbaugh, P. J.; |
| 301 |  | Ul-Hasan, S.; van der Hooft, J. J. J.; Vargas, F.; Vázquez-Baeza, Y.; Vogtmann, E.; von |

| 302 |  | Hippel, M.; Walters, W.; Wan, Y.; Wang, M.; Warren, J.; Weber, K. C.; Williamson, C. |
| --- | --- | --- |
| 303 |  | H. D.; Willis, A. D.; Xu, Z. Z.; Zaneveld, J. R.; Zhang, Y.; Zhu, Q.; Knight, R.; Caporaso, |
| 304 |  | J. G. Reproducible, Interactive, Scalable and Extensible Microbiome Data Science Using |
| 305 |  | QIIME 2. *Nat. Biotechnol.* **2019**, *37* (8), 852–857. <https://doi.org/10.1038/s41587-019-> |
| 306 |  | 0209-9. |
| 307 | (9) | Callahan, B. J.; McMurdie, P. J.; Rosen, M. J.; Han, A. W.; Johnson, A. J. A.; Holmes, S. |
| 308 |  | P. DADA2: High-Resolution Sample Inference from Illumina Amplicon Data. *Nat.* |
| 309 |  | *Methods* **2016**, *13* (7), 581–583. <https://doi.org/10.1038/nmeth.3869>. |
| 310 | (10) | Rognes, T.; Flouri, T.; Nichols, B.; Quince, C.; Mahé, F. VSEARCH: A Versatile Open |
| 311 |  | Source Tool for Metagenomics. *PeerJ* **2016**, *4*, e2584. <https://doi.org/10.7717/peerj.2584>. |
| 312 | (11) | Quast, C.; Pruesse, E.; Yilmaz, P.; Gerken, J.; Schweer, T.; Yarza, P.; Peplies, J.; |
| 313 |  | Glöckner, F. O. The SILVA Ribosomal RNA Gene Database Project: Improved Data |
| 314 |  | Processing and Web-Based Tools. *Nucleic Acids Res.* **2013**, *41* (D1), D590–D596. |
| 315 |  | <https://doi.org/10.1093/nar/gks1219>. |
| 316 | (12) | Göker, M.; Christensen, H.; Fingerle, V.; Kostovski, M.; Margos, G.; Moore, E. R. B.; |
| 317 |  | Oren, A.; Patrick, S.; Reischl, U.; Vázquez-Boland, J. A. List of Recommended Names |
| 318 |  | for Bacteria of Medical Importance: Report of the Ad Hoc Committee on Mitigating |
| 319 |  | Changes in Prokaryotic Nomenclature. *Int. J. Syst. Evol. Microbiol.* **2025**, *75* (10), |
| 320 |  | 006943. <https://doi.org/10.1099/ijsem.0.006943>. |
| 321 | (13) | Cabezas, M. P.; Fonseca, N. A.; Muñoz-Mérida, A. MIMt: A Curated 16S rRNA |
| 322 |  | Reference Database with Less Redundancy and Higher Accuracy at Species-Level |
| 323 |  | Identification. *Environ. Microbiome* **2024**, *19* (1), 88. <https://doi.org/10.1186/s40793-> |
| 324 |  | 024-00634-w. |
| 325 | (14) | Parks, D. H.; Chaumeil, P.-A.; Mussig, A. J.; Rinke, C.; Chuvochina, M.; Hugenholtz, P. |
| 326 |  | GTDB Release 10: A Complete and Systematic Taxonomy for 715 230 Bacterial and |
| 327 |  | 17 245 Archaeal Genomes. *Nucleic Acids Res.* **2026**, *54* (D1), D743–D754. |
| 328 |  | <https://doi.org/10.1093/nar/gkaf1040>. |
| 329 | (15) | Camacho, C.; Coulouris, G.; Avagyan, V.; Ma, N.; Papadopoulos, J.; Bealer, K.; Madden, |
| 330 |  | T. L. BLAST + : Architecture and Applications. **2009**, *9*, 1–9. |
| 331 |  | <https://doi.org/10.1186/1471-2105-10-421>. |
| 332 | (16) | Girard, L.; Lood, C.; Höfte, M.; Vandamme, P.; Rokni-Zadeh, H.; Van Noort, V.; Lavigne, |
| 333 |  | R.; De Mot, R. The Ever-Expanding Pseudomonas Genus: Description of 43 New Species |
| 334 |  | and Partition of the Pseudomonas Putida Group. *Microorganisms* **2021**, *9* (8), 1766. |
| 335 |  | <https://doi.org/10.3390/microorganisms9081766>. |
| 336 | (17) | Rudra, B.; Gupta, R. S. Phylogenomic and Comparative Genomic Analyses of Species of |
| 337 |  | the Family Pseudomonadaceae: Proposals for the Genera Halopseudomonas Gen. Nov. |
| 338 |  | and Atopomonas Gen. Nov., Merger of the Genus Oblitimonas with the Genus |
| 339 |  | Thiopseudomonas, and Transfer of Some Misclassified Species of the Genus |
| 340 |  | Pseudomonas into Other Genera. *Int. J. Syst. Evol. Microbiol.* **2021**, *71* (9), 005011. |
| 341 |  | <https://doi.org/10.1099/ijsem.0.005011>. |
| 342 | (18) | Saati-Santamaría, Z.; Peral-Aranega, E.; Velázquez, E.; Rivas, R.; García-Fraile, P. |
| 343 |  | Phylogenomic Analyses of the Genus Pseudomonas Lead to the Rearrangement of Several |
| 344 |  | Species and the Definition of New Genera. *Biology* **2021**, *10* (8), 782. |
| 345 |  | <https://doi.org/10.3390/biology10080782>. |
| 346 | (19) | Lalucat, J.; Gomila, M.; Mulet, M.; Zaruma, A.; García-Valdés, E. Past, Present and |
| 347 |  | Future of the Boundaries of the Pseudomonas Genus: Proposal of Stutzerimonas Gen. |
| 348 |  | Nov. *Syst. Appl. Microbiol.* **2022**, *45* (1), 126289. |
| 349 |  | <https://doi.org/10.1016/j.syapm.2021.126289>. |
| 350 | (20) | Rudra, B.; Gupta, R. S. Phylogenomics Studies and Molecular Markers Reliably |
| 351 |  | Demarcate Genus Pseudomonas Sensu Stricto and Twelve Other Pseudomonadaceae |

352 Species Clades Representing Novel and Emended Genera. *Front. Microbiol.* **2024**, *14*.
353 <https://doi.org/10.3389/fmicb.2023.1273665>.
354 (21) Pruesse, E.; Peplies, J.; Glöckner, F. O. SINA: Accurate High-Throughput Multiple
355 Sequence Alignment of Ribosomal RNA Genes. *Bioinformatics* **2012**, *28* (14), 1823–
356 1829. <https://doi.org/10.1093/bioinformatics/bts252>.
357 (22) Wong, T. K. F.; Ly-Trong, N.; Ren, H.; Baños, H.; Roger, A. J.; Susko, E.; Bielow, C.;
358 Maio, N. D.; Goldman, N.; Hahn, M. W.; Huttley, G.; Lanfear, R.; Minh, B. Q. IQ-TREE
359 3: Phylogenomic Inference Software Using Complex Evolutionary Models. **2025**.
360 (23) Letunic, I.; Bork, P. Interactive Tree of Life (iTOL) v6: Recent Updates to the
361 Phylogenetic Tree Display and Annotation Tool. *Nucleic Acids Res.* **2024**, *52* (W1), W78–
362 W82. <https://doi.org/10.1093/nar/gkae268>.
363 (24) Lahti,	L.;	Shetty,	S.	Microbiome	R	Package.	**2017**.
364 <https://doi.org/10.18129/B9.bioc.microbiome>.
365 (25) Kassambara, A. Rstatix: Pipe-Friendly Framework for Basic Statistical Tests, 2026.
366 <https://cran.r-project.org/web/packages/rstatix/index.html> (accessed 2026-07-09).
367 (26) Oksanen, J.; Simpson, G. L.; Blanchet, F. G.; Kindt, R.; Legendre, P.; Minchin, P. R.;
368 O’Hara, R. B.; Solymos, P.; Stevens, M. H. H.; Szoecs, E.; Wagner, H.; Barbour, M.;
369 Bedward, M.; Bolker, B.; Borcard, D.; Borman, T.; Carvalho, G.; Chirico, M.; Caceres,
370 M. D.; Durand, S.; Evangelista, H. B. A.; FitzJohn, R.; Friendly, M.; Furneaux, B.;
371 Hannigan, G.; Hill, M. O.; Lahti, L.; Martino, C.; McGlinn, D.; Ouellette, M.-H.; Cunha,
372 E. R.; Smith, T.; Stier, A.; Braak, C. J. F. T.; Weedon, J. Vegan: Community Ecology
373 Package, 2026. <https://cran.r-project.org/web/packages/vegan/index.html> (accessed 2026-
374 04-14).

375	(27) Arbizu, P. M. **2020**.

376 (28) Bartlett, A.; Padfield, D.; Lear, L.; Bendall, R.; Vos, M. A Comprehensive List of
377 Bacterial	Pathogens	Infecting	Humans.	*Microbiology*	**2022**,	*168*	(12).
378 <https://doi.org/10.1099/mic.0.001269>.
379 (29) LeCo project; Gabrielli, M.; Cavallaro, A.; Eichelberg, A.; Margot, C.; Hammes, F.;
380 Kohler, J.; Sigrist, J.; Natasha, R. LeCo Project, LegioSpecies - an Updated and
381 Referenced	List	of	Legionellaceae	Species.	**2024**.
382 <https://doi.org/10.5281/zenodo.11072745>.

383

384

385

### Supplementary Figures

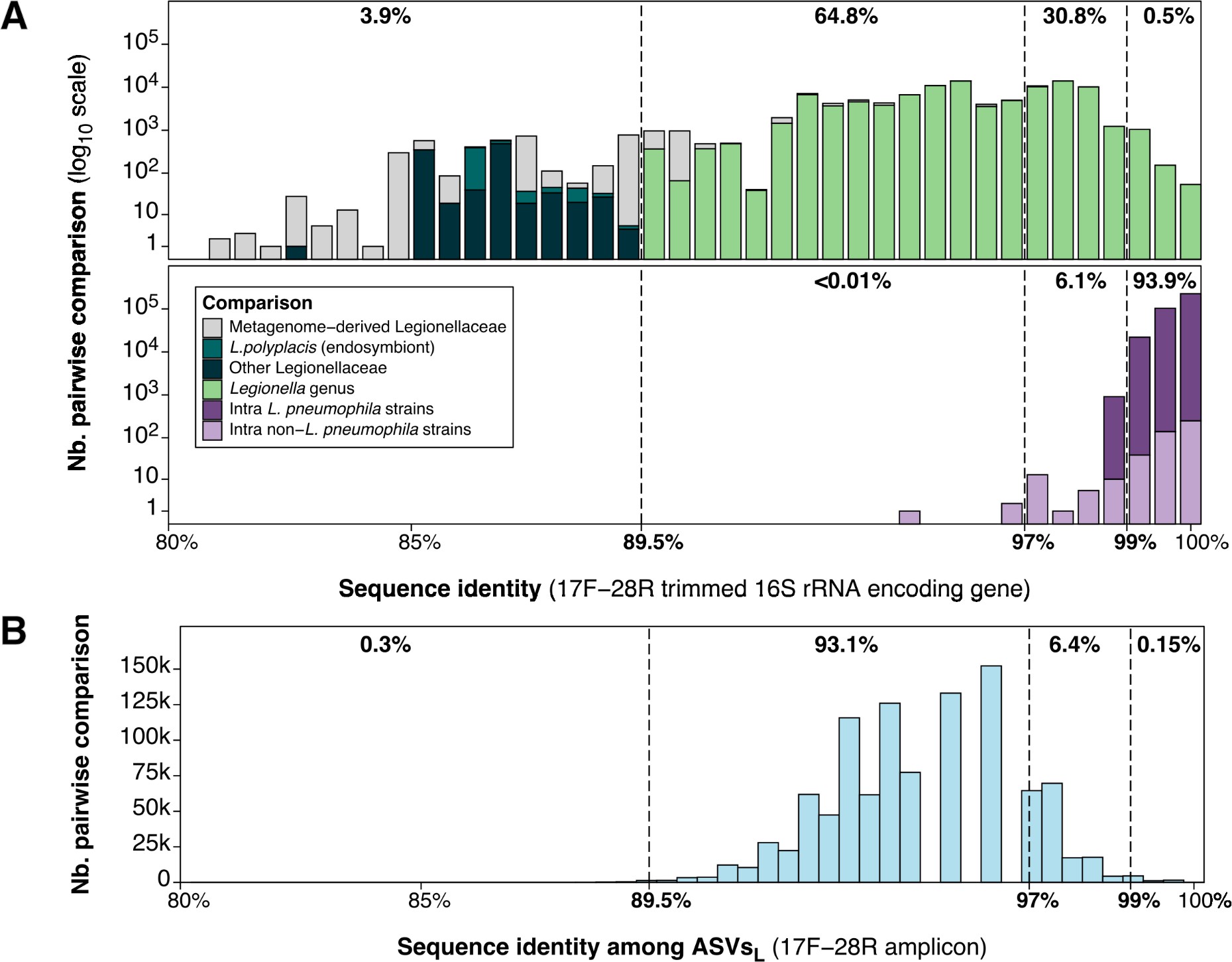

#### Fig. S1: Relatedness among sequences from the *Legionella* curated database and among this study’s ASVsL.

**A**: Inter- and intra-species comparison among *Legionella* strains 16S rRNA encoding gene trimmed with the 17F-28R primers (1235 sequences). **B**: Sequence comparisons among ASVsL. The intra- and inter-species sequence identity thresholds are indicated in bold: distantly related (<97%), closely related (95-99%) and very closely related (>99%).

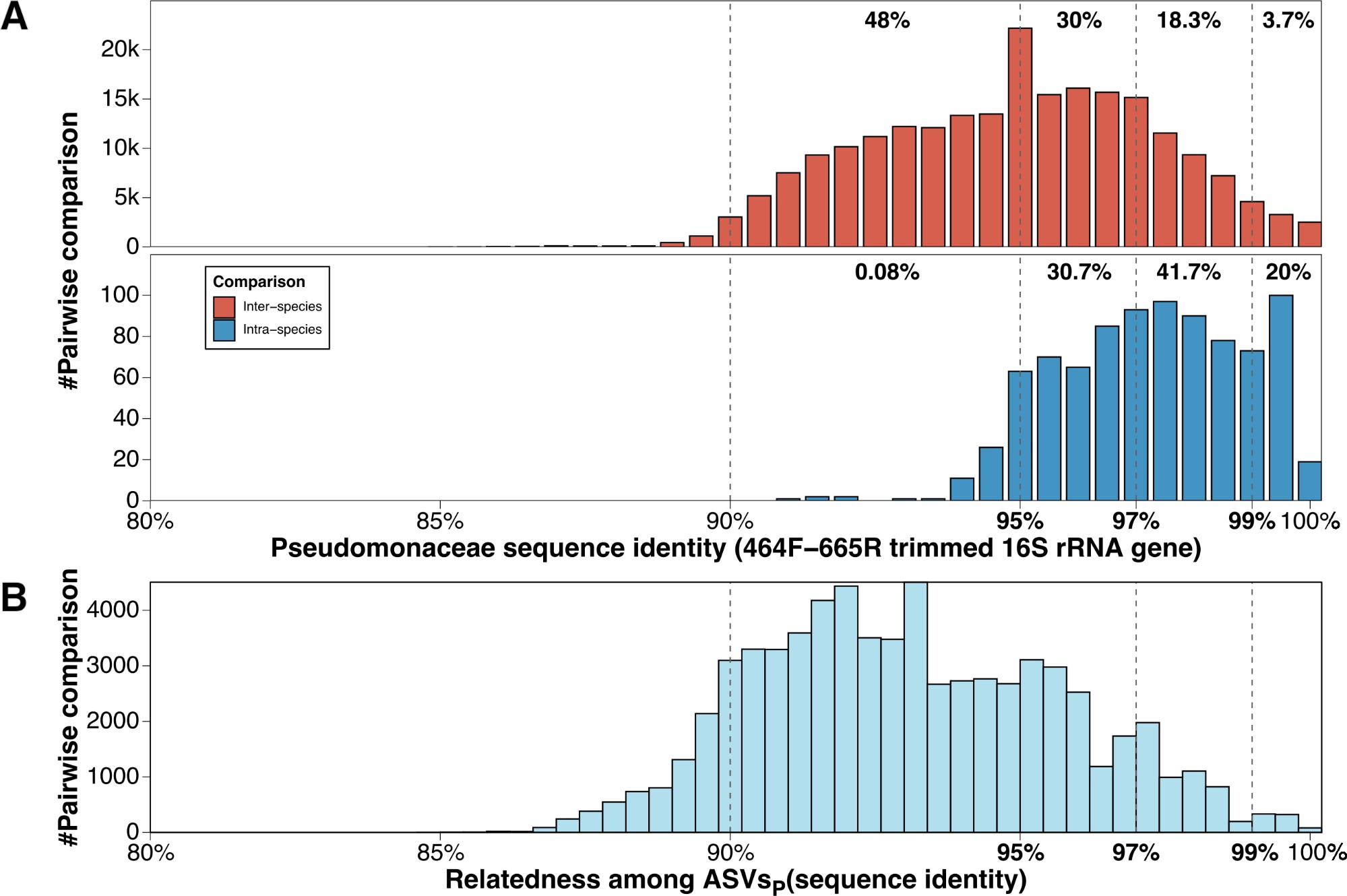

#### Fig. S2: Relatedness among sequences from the Pseudomonadaceae curated database and among this study’s ASVsP.

**A**: Inter- and intra-species comparison among Pseudomonadaceae strains 16S rRNA encoding gene trimmed with the 464F-665R primers (734 sequences). **B**: Sequence comparisons among ASVsP. The intra- and inter-species sequence identity thresholds are indicated in bold: distantly related (<95%), closely related (95-99%) and very closely related (>99%).

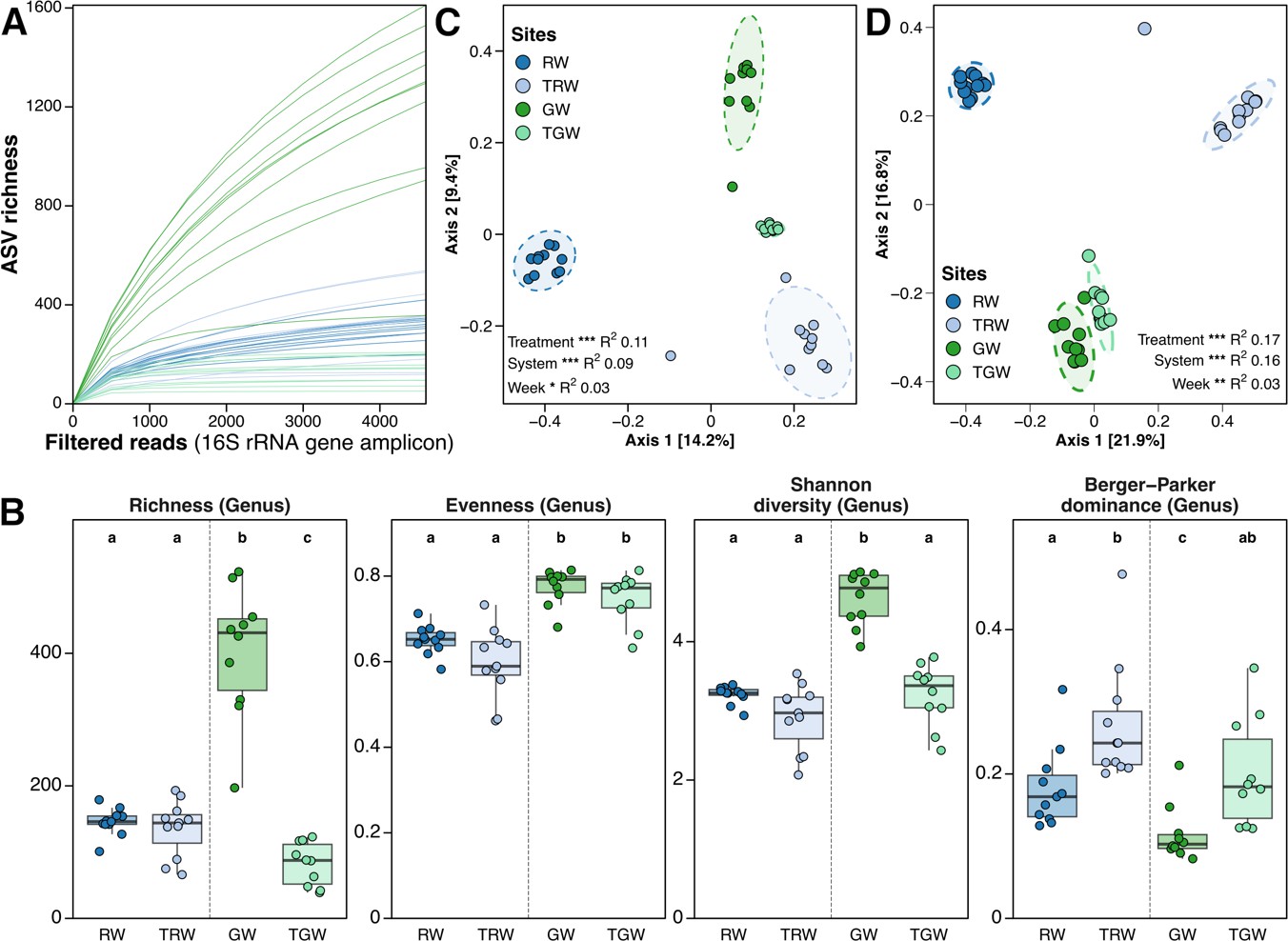

#### Fig. S3: Additional prokaryotic community diversity and composition among water sources and their respective treated waters.

**A**: Rarefaction curve at the ASV level (set at 4600 reads). **B**: Alpha-diversity indices based on prokaryotic genera (Richness, Evenness, Shannon diversity and Berger-Parker dominance). Letters refer to statistical differences using Kruskall-Wallis and Wilcoxon *post-hoc* tests (Bonferroni adjusted *p*-values). **C-D**: PCoA plot (Jaccard (**C**) and Bray-Curtis (**D**) dissimilarity) based on prokaryotic ASVs. Ellipses represent 95% confidence intervals and permanova statistics are displayed (Treatment: raw *vs*. treated water; System: river (raw and treated) *vs*. groundwater (raw and treated); Week:1-12). Stars refer to statistical significance (* *p*<0.05, ** *p*<0.01 and *** *p*<0.001).

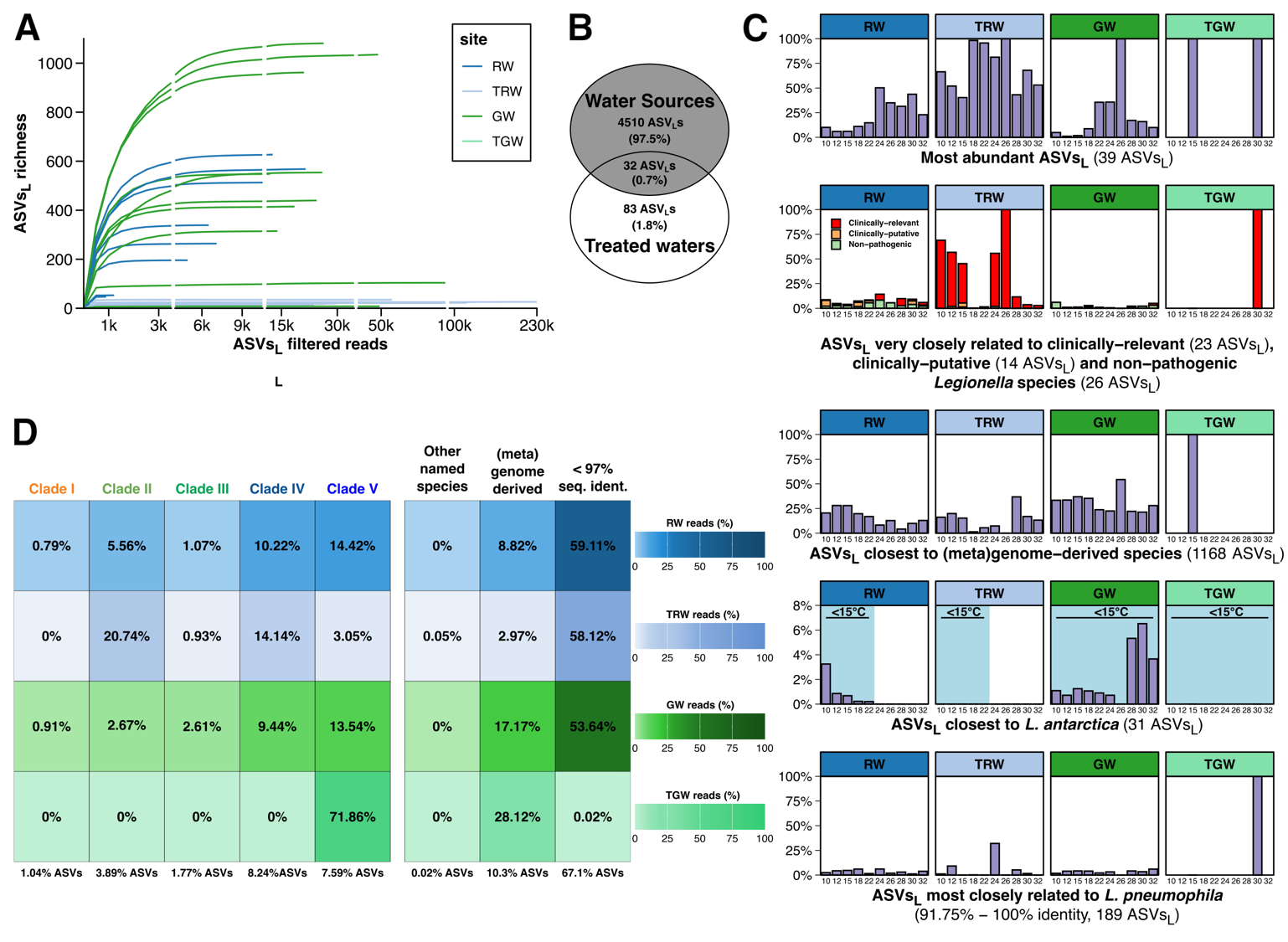

#### Fig. S4: Additional *Legionella* community diversity and composition among water sources and their respective treated waters.

**A:** Rarefaction curves based on *Legionella* filtered reads for each sample. **B**: Venn diagram based on ASVsL. **C**: Summed relative abundance of most abundant ASVsL (see Methods), ASVsL closest to (meta)genome-derived *Legionella* or legionellaceae species, ASVsL very closely related to clinically- and non clinically-relevant *Legionella* species and ASVsL most closely related to *L. antarctica* and *L. pneumophila*. **D**: Relative abundance and number (below, in parenthesis) of ASVsL according to their phylogenomic clade identified from the *Legionella* core-genome phylogeny (Gomez-Valero *et al.* 2019) at each site. ASVsL affiliated to species without known *Legionella* cluster affiliation, to (meta)genome-derived species or affiliated at less that 97% seq. identity were displayed apart.

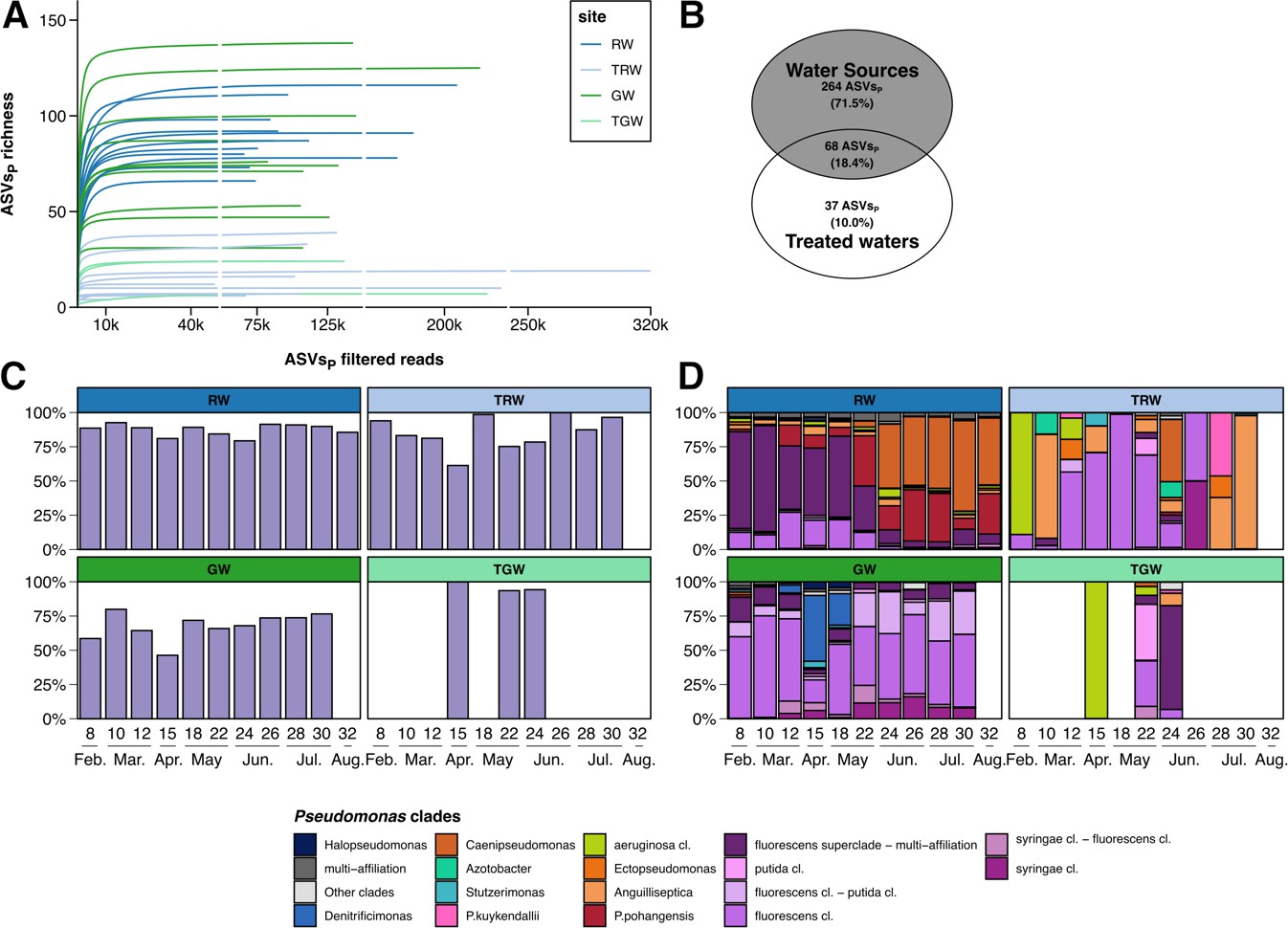

#### Fig. S5: Additional *Pseudomonas* community diversity and composition among water sources and their respective treated waters.

**A:** Rarefaction curves based on *Pseudomonas* filtered reads for each sample. **B**: Venn diagram based on ASVsP. **C**: Summed relative abundance of most abundant ASVsP (see Methods). **D**: Relative abundance of most abundant *Pseudomonas* clades at the species if unique affiliation was possible, genera or clade within the *P. fluorescens* superclade (Rudra and Gupta 2024).

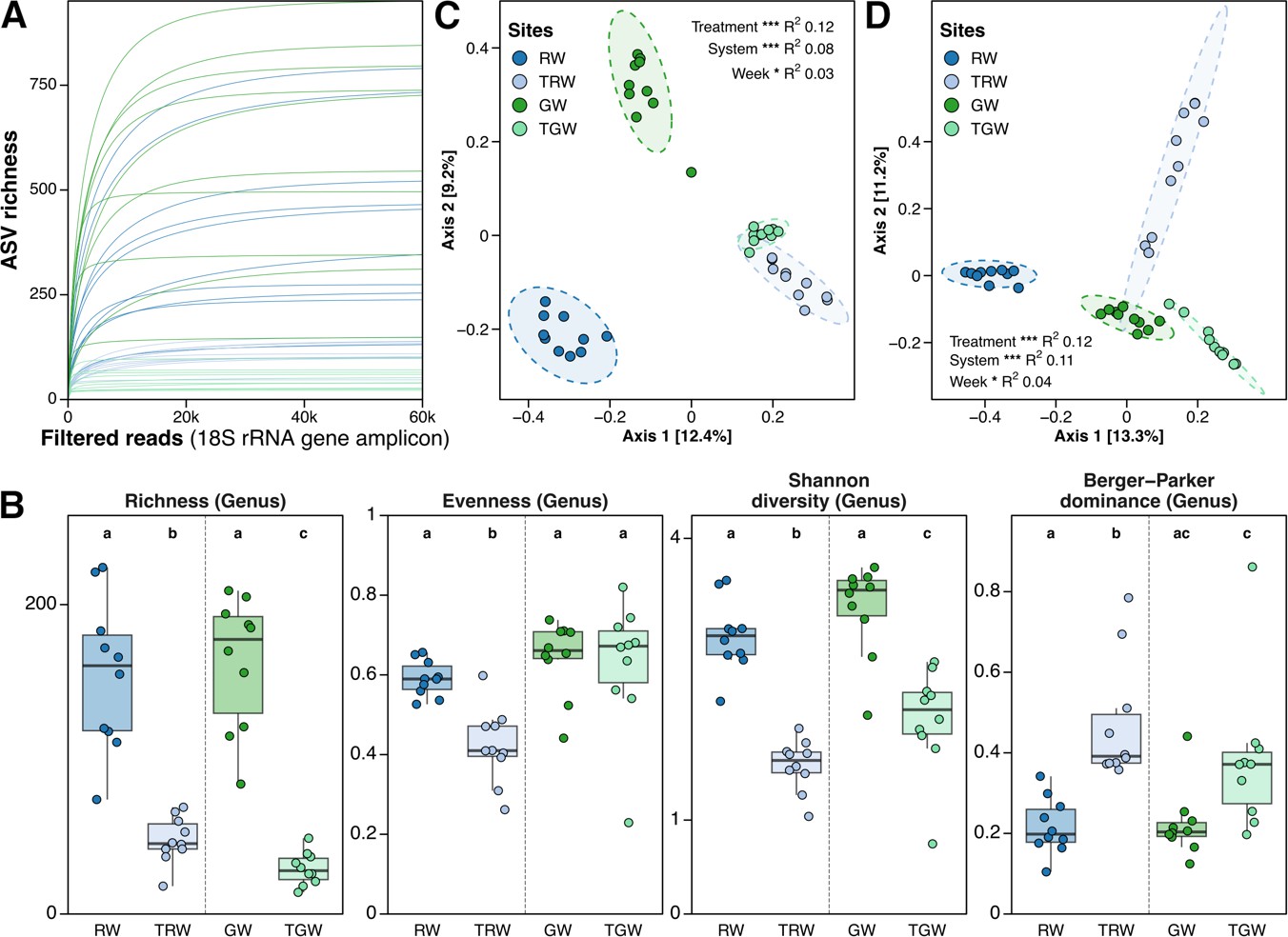

#### Fig. S6: Additional eukaryotic diversity and community composition

**A**: Rarefaction curve at the ASV level (set at 60,000 reads). **B**: Alpha-diversity indices based on eukaryotic genera (Richness, Evenness, Shannon diversity and Berger-Parker Shannon diversity). Letters refer to statistical differences using Kruskall-Wallis and Wilcoxon *post-hoc* tests (Bonferroni adjusted *p*-values). **C-D**: PCoA plot (Jaccard (**C**) and Bray-Curtis (**D**) dissimilarity) based on eukaryotic ASVs. Ellipses represent 95% confidence intervals and permanova statistics are displayed (Treatment: raw *vs*. treated water; System: river (raw and treated) *vs*. groundwater (raw and treated); Week:8-30). Stars refer to statistical significance (* *p*<0.05, ** *p*<0.01 and *** *p*<0.001).
